# Mechanical, Biochemical, and Multicellular Effects on Vessel Network Morphometrics in a Microfluidic Vasculature-on-a-Chip

**DOI:** 10.1101/2025.08.28.672831

**Authors:** Han Shao, Edmond W. K. Young

## Abstract

Microvascular networks (MVNs) formed via endothelial cell self-assembly in 3D hydrogels have emerged as a widely used platform for modeling vascularized tissues and studying vascular pathophysiology. Conventional MVN systems incorporate supporting fibroblasts and may include biochemical cues such as VEGF, FGF, or S1P, as well as mechanical stimuli like luminal flow, yet the impact of these variables on MVN morphology and function remains incompletely understood. Here, we systematically investigated the effects of fibroblast concentration, fibroblast-conditioned media, angiogenic factors, and luminal flow on the morphology, perfusability, and vessel wall integrity of MVNs cultured in a microfluidic vasculature-on-a-chip. In addition to standard branch-based metrics such as vessel coverage area and diameter, we developed and applied novel void-based morphological parameters that quantify the size, shape, and distribution of vessel-free spaces to capture subtle differences across MVN culture conditions. Our results demonstrate that high fibroblast-to-endothelial cell ratios accelerate MVN formation but promote excessive vessel fusion, while MVNs cultured without fibroblasts—using only conditioned media or soluble factors—exhibited patch-like, non-physiological morphology with reduced branch formation. Direct inclusion of fibroblasts proved essential for promoting the thin, interconnected vascular structures characteristic of in vivo microvasculature and could not be substituted by soluble cues alone. Furthermore, the presence or absence of fibroblasts modulated MVN responsiveness to luminal flow. Overall, our void-based analysis method enabled more sensitive discrimination of MVN morphological features than traditional branch-based metrics and offers a reduced-data, high-content approach suitable for integration with machine learning and AI-assisted image analysis pipelines. This platform provides a new framework for optimizing MVN culture protocols and advancing vascular tissue engineering studies, particularly for the advancement of organ-on-a-chip (OOC) and microphysiological systems.

## Introduction

Blood vessels deliver essential nutrients and oxygen necessary for tissue viability. Without vascularization, tissues cannot grow beyond the maximum oxygen diffusion distance of ∼200 μm [1]. Since functional blood vessels are critical for maintaining normal organ physiology, understanding the role of vasculature in human health and disease is crucial for elucidating disease mechanisms such as tumor growth and metastasis, developing treatments for vascular-related disorders, and advancing tissue engineering strategies that require functional vascular networks to support the survival and integration of engineered tissues. Experimental models of the vasculature are thus critical to uncovering mechanisms of vascular function, disease progression, and therapeutic response.

Conventional *in vivo* animal models and *in vitro* models such as static cell cultures have been invaluable for advancing our understanding of human biology and disease mechanisms associated with the vasculature; however, they possess significant limitations, including poor physiological relevance and limited control over the cellular microenvironment in the case of static cultures, and lack of human-specific responses, high cost, and ethical concerns in the case of animal models. As a result, bioengineered tissue models using microfluidic devices – commonly known as microphysiological (MPS) or organ-on-chip (OOC) systems – are becoming increasingly popular for organ and disease modeling. These OOC platforms offer the advantages of enabling precise fluidic control, reducing reagent consumption, and providing precise recapitulation of tissue-and organ-level organization [2,3]. By further incorporating human-relevant cells into these systems, OOCs provide accurate mimicry of human biology while avoiding translational challenges related to interspecies differences [4].

To date, vascularized OOC models have been established for many organ systems, including the lung[5], heart[6,7], and brain[8]. These models are used for myriad applications including to study cardiovascular diseases such as atherosclerosis and thrombosis [5], respiratory conditions like pulmonary edema [9] and neurological diseases such as blood-brain barrier dysfunction[10] and neuroinflammation [11]. In addition to these applications, vascularized OOC platforms are utilized in cancer research, including studies of tumour angiogenesis [12,13], cancer cell invasion and intravasation[14], extravasation[15], tumour-immune interaction[16], and the process of intussusceptive angiogenesis[17]. Further developments and application of vascularized OOC systems have been summarized in recent reviews[18–21]. In terms of design formats, the on-chip vascular components within these OOC systems are typically categorized into one of these three types: 1) vessel barriers formed on flat surfaces [5,22,23], 2) single tubular vessel lumens [24–27], and 3) networks of interconnected vessel branches[28,6]. In relation to vessel networks, two main fabrication approaches are typically employed: 1) lining prefabricated network structures with endothelial cells [29–31] and 2) inducing vessel network formation within 3D hydrogels through angiogenesis[32,33] or vasculogenesis via self-assembly [34–43]. The latter are generally referred to as self-assembled microvascular networks (MVNs).

MVNs represent one of the most physiologically relevant *in vitro* vascular models. Typically, MVNs are cultured in microfluidic channels where endothelial cells self-assemble within a fibrin gel matrix, often supported by co-cultured fibroblasts and supplemented with angiogenic growth factors such as VEGF[44] and mechanical cues like perfusion flow [25,45,46]) [47]. However, fibroblasts in these systems tend to proliferate rapidly and continuously, which can obstruct microfluidic channels, disrupt vessel perfusion, and ultimately lead to vessel lumen collapse during long-term culture. Therefore, strategies that reduce or replace fibroblasts could enhance the stability and longevity of MVN cultures.

MVNs typically form within approximately one week, with total culture durations usually under two weeks; only a few studies have reported maintaining viable networks for up to a month[48]. Unlike the relatively stable microvasculature found *in vivo*, *in vitro* MVNs often undergo dynamic morphological changes both during and after network formation. Understanding these baseline morphological shifts is critical to accurately assess the impact of additional treatments or disease models without confounding effects from inherent MVN remodeling. However, such temporal dynamics are frequently overlooked in existing analyses.

A wide variety of MVN culture protocols have been reported, resulting in networks with highly variable morphologies—from irregular, short, and wide vessels with inconsistent diameters to more physiological networks consisting of thin, elongated branches. Current analysis tools, such as AngioTool[49], along with common metrics (e.g., number of branches, total branch length), are optimized for *in vivo*-like vasculature with slender, lengthy structures. Consequently, these tools fail to capture many subtle but important features unique to *in vitro* MVNs, leading to incomplete or inaccurate morphological assessments.

In this study, we established microfluidic MVNs on-chip and systematically tested the effects of varying fibroblast ratios, biochemical factors in the culture medium, and mechanical flow conditions on MVN formation and growth dynamics. Our objectives were twofold: 1) to explore alternatives to fibroblast co-culture for improving the stability of long-term MVN cultures, and 2) to examine how different culture protocols influence MVN morphology. To capture morphological characteristics that are often overlooked in the literature, we introduced novel image analysis parameters, including vessel network void space number, roundness, and alignment angle—features that reflect vessel density, shape, and directional organization. Using a newly developed image analysis pipeline, we comprehensively quantified these parameters across various MVN culture conditions. This study presents a new framework for assessing MVN morphology and growth dynamics, offering deeper insight into how culture variables influence microvascular network formation *in vitro*.

## Materials and Methods

### Cell Culture

Red fluorescent protein-expressing human umbilical vein endothelial cells (RFP-HUVECs, cAP-0001RFP) and green fluorescent protein-expressing human adult lung fibroblasts (GFP-hLFs, cAP-0033GFP) were purchased from Angio-Proteomie (Boston, MA, USA) and cultured on T75 flasks coated with Quick Coating solution (cAP-01, Angio-Proteomie) according to manufacturer’s protocol. RFP-HUVECs were cultured in EGM-2 media with EGM-2 BulletKit supplements (CC-3162, Lonza) and used between passages 6 to 7. GFP-hLFs were cultured in Dulbecco’s Modified Eagle Medium (DMEM, 11885-084, Gibco, Thermo Fisher Scientific) supplemented with 10% fetal bovine serum (FBS, 12483-020, Gibco) and used between passage 7 to 9. All cells were cultured in incubator at 37°C and 5% CO_2_. Media was changed every other day, and cells were passaged when confluency reached ∼ 90%.

### Mold Fabrication

Positive-relief molds containing the MVN-on-a-chip design were fabricated from poly(methylmethacrylate) (PMMA) sheets (McMaster-Carr, IL, USA) by micromilling. The mold model and machine toolpaths were generated in Autodesk Fusion360 and then translated into G-code for micromilling (Tormach PCNC770, Waunakee, WI, USA). To remove roughness of the milled surface, the finished PMMA mold was subjected to a chloroform vapor polishing treatment. Chloroform (366927-1L, Sigma-Aldrich) was added into a glass petri dish that was subsequently placed in a container filled with a thin layer of water. The PMMA mold was taped to the bottom of a second petri dish with the milled surface facing up. The second petri dish was flipped upside down and placed above the chloroform-containing petri dish, thus exposing the milled surface to chloroform vapor. Note that the edge of the PMMA-containing petri dish was submerged in the water of the external container to create a sealed vapor chamber. The PMMA mold was exposed to the chloroform for two 5-minute periods at room temperature, separated by 10 minutes at room temperature away from the chloroform. The chloroform-treated PMMA mold was then kept at room temperature for a day to allow the PMMA to re-anneal via polymer chain reorganization. To further remove surface contaminants of vapor-polished PMMA mold, a sacrificial layer of poly(dimethylsiloxane) (PDMS) was poured on top, allowed to cure, and then peeled away to reveal a smoothed polished PMMA surface ready for PDMS device fabrication.

### PDMS Device Fabrication

PDMS (Sylgard 184) silicone elastomer base and curing agent was mixed at a 10:1 ratio, respectively, and poured over the clean and polished PMMA mold. The PDMS was then cured in an oven at 37°C overnight to minimize PDMS shrinkage that typically occurs at elevated baking temperatures. Device reservoirs were made with a 6-mm diameter biopsy punch. Cured PDMS casts and glass slides were cleaned in an ultrasonic bath while first submerged in 70% ethanol for 5 min, then subsequently submerged in deionized water for another 5 min, and then blow-dried with compressed air. The glass and PDMS surfaces were then treated with air plasma in a plasma chamber for 2.5 min, immediately pressed together and then baked on a hotplate at 80°C for 5 min to complete the bonding step.

### Device Preparation Prior to Cell Seeding

One day before cell seeding, the PDMS-glass microfluidic device was internally filled with 70% ethanol in all its channels, further disinfected externally by soaking in 70% ethanol for 40 min, and then flushed internally with Dulbecco’s Phosphate Buffered Saline (PBS). To prime the microfluidic device, EGM-2 media (CC-3162, Lonza) was loaded into the device reservoirs and all microchannels and then kept in an incubator at 37°C overnight. On the day of cell seeding, the remaining EGM-2 in the device was aspirated and the device flushed with PBS again. To improve gel attachment to interior microchannel walls, the microchannels were coated with 10 mg/mL human fibronectin (F0895, Sigma-Aldrich) in an incubator at 37°C for one hour.

### Device Seeding with Fibroblasts

The cocultured MVN was formed by culturing RFP-HUVECs and GFP-hLFs together in a fibrin gel. The final fibrin gel solution consisted of 4 mg/mL bovine fibrinogen (F8630, Sigma-Aldrich), 1 U/mL bovine thrombin (T4648, Sigma-Aldrich) and 0.2 mg/mL type I collagen (354249, Corning) diluted in EGM-2 supplemented with 1% (v/v) Penicillin-Streptomycin (P/S; Gibco). Cells were trypsinized and resuspended in the gel precursor mixture to produce a final concentration of (a) 5×10^6^ cells/mL for both RFP-HUVECs and GFP-hLFs for a 1:1 EC:FB ratio, or (b) 5×10^6^ cells/mL RFP-HUVEC and 1×10^6^ cells/mL GFP-hLFs for a 5:1 EC:FB ratio. When making the mixture of cells in gel, cells were resuspended in a partial gel solution containing only thrombin and collagen. Fibrinogen was added as the last step, immediately before loading the complete mixture of cells in gel into the middle vessel chamber of the device. After loading, the device was placed in incubator at 37°C for 10 min to polymerize the gel. Then, EGM-2 supplemented with 1% (v/v) P/S and 0.05 TIU/mL aprotinin (A3428, Sigma Aldrich) was added into the two side channels and the device was incubated at 37°C for an additional 15 min.

To create an endothelial monolayer in the side channels, a separate flask of RFP-HUVECs was trypsinized and resuspended in EGM-2 supplemented with 1% (v/v) P/S at 5×10^6^ cells/mL. The suspended cells were first loaded into the left channels of the device. The device was placed in the incubator for 10 min and tilted at ∼45° (with left channel above right channel) such that the RFP-HUVECs settled onto the left-side fibrin gel interface. Without removing the cell suspension in the left channel, the same seeding procedure was repeated for the right channel with ∼45° tilt in the opposite direction (with right channel above left channel) such that the RFP-HUVECs settled onto the right-side fibrin gel interface. The device was then incubated while lying flat for 10 min to allow RFP-HUVECs to strengthen their attachment. Excess cells were removed by slowly emptying out all remaining liquid from both side channels with a pipette. Each reservoir was filled with 100 μL of EGM-2 supplemented with 1% (v/v) P/S and 0.05 TIU/mL aprotinin. The device was placed in an Omnitray with sacrificial droplets of PBS around the device to reduce media evaporation. MVNs were cultured for 7 days with daily media changes. See Figure S1 for experimental timelines of seeding, media changes, and sample collection.

### Device Seeding without Fibroblasts

All MVNs cultured without fibroblasts followed the same seeding procedure, except the fibroblasts were not added to the mixture of cells in gel. Following the same seeding procedure as described above, RFP-HUVECs were resuspended in the fibrin gel mixture and loaded into the middle channel of the device at 5×10^6^ cells/mL. Endothelial monolayers were then seeded into the two side channels and onto the gel interfaces as described above. See Figure S1 for experimental timelines of seeding, media changes, and sample collection.

### Fibroblast Conditioned Media

Fibroblast conditioned media (FB-CM) was collected from fibroblasts cultured alone (in monoculture) in fibrin gel in the microfluidic device. By replicating the same 3D fibrin gel and microfluidic device environment, we aimed to produce a soluble factor profile that was more representative of the soluble factors secreted by fibroblasts in our FB:EC coculture experiments. Following the same seeding procedure as describe above for cocultures, GFP-hLFs were resuspended in the fibrin gel mixture and loaded into the middle channel of the device at 1×10^7^ cells/mL. After gel polymerization, each reservoir was filled with 100 μL of EGM-2 supplemented with 1% (v/v) P/S and 0.05 TIU/mL aprotinin and placed in the incubator at 37°C. The conditioned media was collected after 24 h, after which 100 μL of fresh EGM-2 supplemented with 1% (v/v) P/S was added to replenish the reservoirs, cultured for a subsequent 24 h, and then collected as conditioned media the next day. All media samples collected from fibroblast monocultures belonging to the same day of culture were pooled together and then supplemented with additional 2.5 mg/mL aprotinin before use as our FB-CM in subsequent MVN-on-a-chip experiments.

For experiments using FB-CM, the same procedure as described in *Device Seeding without Fibroblasts* was used. On the day of seeding (Day 0), MVNs with ECs only were incubated with EGM-2 supplemented with 1% (v/v) P/S and 10 mg/mL aprotinin in the reservoirs. Fibroblast monocultures were also seeded on the same day so that FB-CM can be collected the next day. From Day 1 to Day 6, FB-CM were collected from the fibroblast monoculture devices each day, pooled together, supplemented with 0.0125 TIU/mL aprotinin, and then loaded into the EC-only MVN devices. The resulting MVNs were those cultured with FB-CM as a replacement for fibroblasts. See Figure S1 for experimental timelines of seeding, media changes, and sample collection.

### Angiogenic Factors

Three angiogenic factors were added endogenously to induce vessel network formation: human VEGF165 (293-VE, Biotechne), bovine basic FGF (bFGF, CYT-264, Prospec), and Sphingosine-1-Phosphate (S1P, S9666, Sigma Aldrich). VEGF165 and bFGF were reconstituted according to manufacturer’s protocol. S1P was diluted in a 1 mg/mL solution of bovine serum albumin (BSA, A7906, Sigma Aldrich). The three factors were added into the culture media at different concentrations for the various conditions according to Table 1. Note that the EGM-2 BulletKit includes 2 ng/mL VEGF and 4 ng/mL FGF of its own native supplements. For the conditions in which native VEGF and FGF are not listed, the VEGF and FGF from the BulletKit were not added when making the supplemented media. All vessel network cultures involving endogenously added angiogenic factors used the procedure described in *Device Seeding without Fibroblasts.* Angiogenic factors were added to the EGM-2 media with 1% (v/v) P/S and 0.05 TIU/mL aprotinin according to the timing and the concentrations listed in Table 1 (also see Figure S1). For the last 4 conditions which investigate the effect of different VEGF concentrations, VEGF and FGF from the EGM-2 supplement kit was not added. The same angiogenic factor concentrations were used for Day 0 to Day 2. Different VEGF165 concentrations was introduced from Day 3 to Day 7.

**Table 1:** Experimental Conditions and Labels with Angiogenic Factor Concentrations.

| *Condition Labels* | *Specific Conditions* | *VEGF (ng/mL)* | *FGF (ng/mL)* | *S1P (nmol/mL)* | *Flow* | *Fibroblast involved?* |
| --- | --- | --- | --- | --- | --- | --- |
| FB5:EC5 | FB = $5 \times 10^6$ cells/mL;<br>EC = $5 \times 10^6$ cells/mL | 2 (from media) | 4 (from media) | 0 | - | Coculture |
| FB1:EC5 | FB = $1 \times 10^6$ cells/mL;<br>EC = $5 \times 10^6$ cells/mL | 2 (from media) | 4 (from media) | 0 | - | Coculture |
| FB1:EC5-flow | FB = $1 \times 10^6$ cells/mL;<br>EC = $5 \times 10^6$ cells/mL;<br>with flow | 2 (from media) | 4 (from media) | 0 | Days 3-7 | Coculture |
| EGM-2 | EGM-2 media | 2 (from media) | 4 (from media) | 0 | - | n/a |
| FB-CM | Fibroblast conditioned media | 2 (from media) | 4 (from media) | 0 | - | CM only |
| Ang-V50 | EGM-2 plus three angiogenic factors (50 ng/mL VEGF165) | 2 (from media) + 50 VEGF165 added | 4 (from media) + 50 bFGF added | 500 | - | n/a |
| Ang-V20 | EGM-2 plus three angiogenic factors (20 ng/mL VEGF165) | 2 (from media) + 20 VEGF165 added | 4 (from media) + 50 bFGF added | 500 | - | n/a |
| Ang-V20-flow | EGM-2 plus three angiogenic factors (50 ng/mL VEGF165) plus flow | 2 (from media) + 20 VEGF165 added | 4 (from media) + 50 bFGF added | 500 | Days 3-7 | n/a |
| VEGF0 | 20 ng/mL VEGF on Day 0-2; switch to VEGF0 Day 3-7 (excludes VEGF and FGF from media) | Day 0-2: 20<br>Day 3-7: 0 | 50 bFGF added | 500 | - | n/a |
| VEGF10 | 20 ng/mL VEGF, on Day 0-2; switch to VEGF10 Day 3-7 (excludes VEGF and FGF from media) | Day 0-2: 20<br>Day 3-7: 10 | 50 bFGF added | 500 | - | n/a |
| VEGF50 | 20 ng/mL VEGF, on Day 0-2; switch to VEGF50 Day 3-7 (excludes VEGF and FGF from media) | Day 0-2: 20<br>Day 3-7: 50 | 50 bFGF added | 500 | - | n/a |
| VEGF100 | 20 ng/mL VEGF, on Day 0-2; switch to VEGF100 Day 3-7 (excludes VEGF and FGF from media) | Day 0-2: 20<br>Day 3-7: 100 | 50 bFGF added | 500 | - | n/a |

### Gravity-Driven Flow

The effect of flow on MVN morphology was studied by establishing gravity-driven flow on the MVN-on-a-chip. Hydrostatic pressure of 5 mmH2O was established beginning on Day 3 after preliminary vessel lumen formation. This pressure was established by creating a media volume difference of ∼282 μL between the inlet and outlet reservoirs of the middle channel. The total volume of media for each device was 400 μL, equivalent to the media volume of our no-flow controls. Gravity-driven flow was re-established every day during media changes.

### Perfusion of Microbeads

Fluorescent microspheres were perfused through the MVN to demonstrate and visualize the perfusability of the MVNs. Blue fluorescent microspheres (Fluospheres, F8829, Thermo Fisher Scientific) were diluted to 1 μL/mL in PBS. To establish the flow of microspheres in the MVN, 100 μL of PBS was added into the two right-hand-side reservoirs, and then 200 μL of the microsphere solution was added into the left-hand-side reservoirs.

### Image Acquisition and Analysis

All images of the MVNs were acquired on an Olympus IX83 inverted microscope with a 4x objective (UPlanFL N 4x/0.13, Olympus). Image analyses were conducted using ImageJ. Data analyses were performed in MATLAB. Detailed image processing steps and the definitions of morphological metrics are included in the *Supplemental Materials*.

### Statistical Analysis

Statistical analyses were performed in SPSS. One-way ANOVA was performed on the data for vessel branch diameters. Post hoc analyses were performed with the Tukey’s HSD test. Mixed ANOVA with repeated measures was performed on the data for percent vessel area, void count, void roundness and void area. Bonferroni correction was applied for simple main effect pairwise comparisons.

## Results and Discussion

### Device and Experimental Design

Our MVN-on-a-chip consists of a PDMS microchannel layer bonded to an underlying glass substrate (Figure 1a). The device configuration consists of three parallel microchannels (Figure 1b): the central microchannel accommodates a 3D hydrogel for supporting a perfusable vessel network (Figure 1c) and the two adjacent side microchannels are lined with a curved monolayer of ECs and are responsible for providing the central microchannel with access to the media in the reservoirs. Phaseguides extend along the length of neighboring microchannels to pin and confine the hydrogel solution within the central microchannel by surface tension [50–52] (Figure 1e). A larger gel reservoir under the gel loading port serves as a buffer for gel loading so that the initial pressure from pipetting is reduced to lower the risk of the gel seeping into the side channels. This gel port design increased the success rate of gel loading. The two side channels are designed to be lined with EC monolayers that anastomose with the formed vessel network in the central channel, thereby allowing network perfusion (Figure 1c). Because phaseguides were employed, the vessel network can be connected to the EC monolayer along the entire channel length, which is distinct from vessel networks using microposts [6,34] or connection ports [53] where vessel networks can only connect to the EC monolayer at defined locations along the channel length.

**Figure 1:**
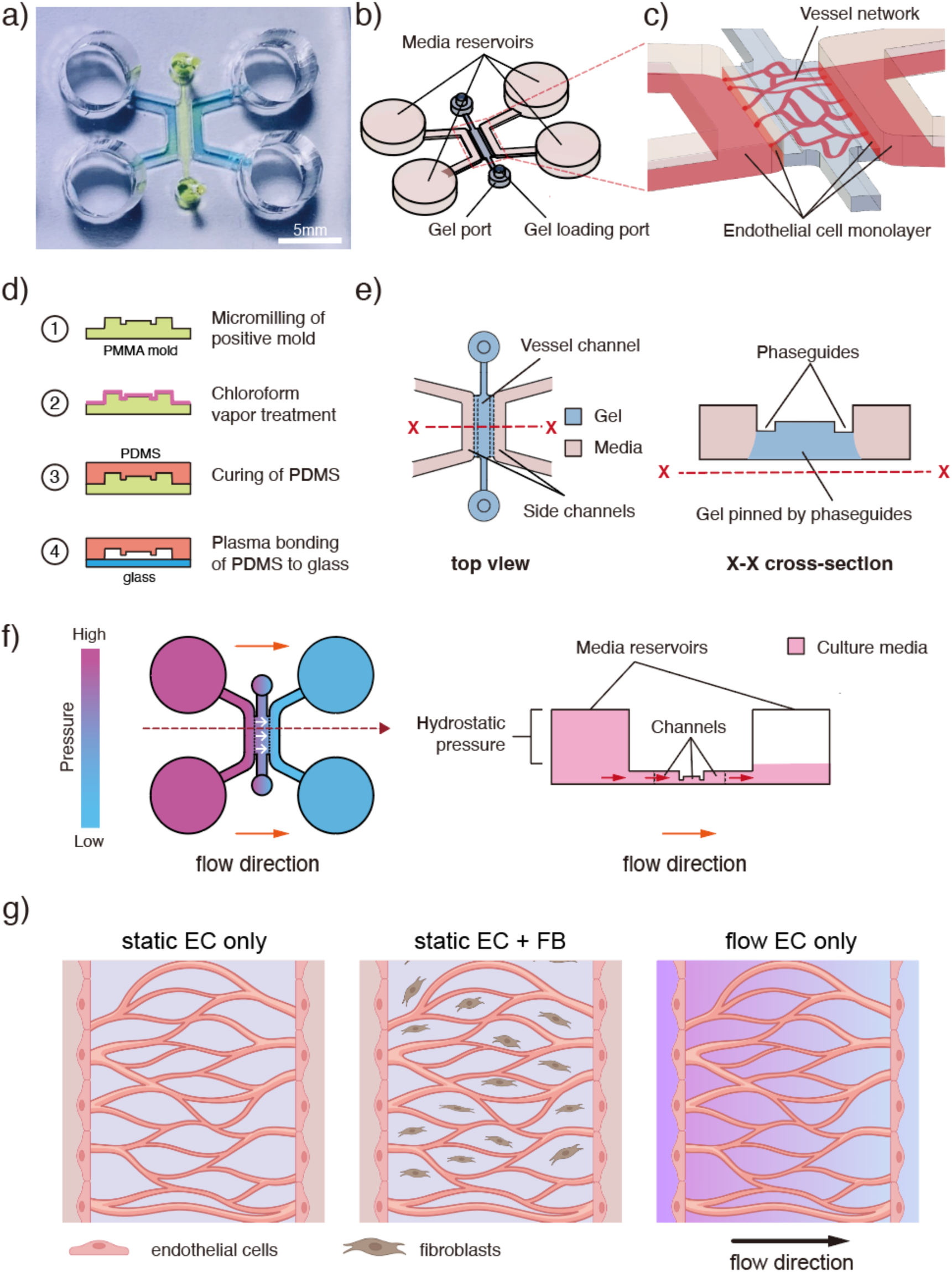
a) Photo of MVN-on-a-chip: blue dye = side channels; yellow dye = middle MVN channel. Scale bar = 5mm. Dimensions in microns (h x w): middle channel 150 x 800; phaseguides 30 x 200; side channels 600 x 800. b), c) and e): Device design. MVN-on-a-chip device consists of a middle channel housing the vessel network in 3D hydrogel and 2 side channels lined with endothelial cells that connect to the vessel network. Channels are separated by phaseguides that pin the gel in the center channel by interfacial tension. d) Device fabrication. f) Gravity flow can be established by hydrostatic pressure of culture media in the reservoirs. g) The device can support a wide range of culture with different cell types and flow condition.

To fabricate the device (Figure 1d), we (i) created a positive-relief micromold of the channel design in PMMA by micromilling, (ii) performed a chloroform vapour treatment on the PMMA micromold, (iii) cured PDMS on the PMMA micromold by soft lithography, and (iv) plasma-bonded the negative-relief PDMS channel layer to glass, as previously described [54]. We chose micromilling for mold fabrication over traditional photolithography with photoresist because it enabled rapid iterations of channel dimensions with high z-direction precision. Micromilling, however, left tool marks on the mold surface that transferred to the PDMS layers, causing surface roughness on the PDMS that lowered its bond strength on glass. To overcome this issue, the PMMA micromold was treated with chloroform vapour to chemically “polish” the tool marks [55,56]. PDMS was cured at low heating temperature to reduce its shrinkage after cooling.

The MVN-on-a-chip can be used to test a variety of microenvironments including vessel networks created from ECs in either monoculture or in cocultures with supporting cells, or under static or flow conditions (Figure 1f and 1g). Of interest in this study were the effects of fibroblast concentration, VEGF concentration, and interstitial flow on the temporal dynamics of MVN morphology (Figure S1). In particular, we surveyed five different culture conditions (Table 1) with a specific goal of determining whether fibroblasts can be replaced by fibroblast conditioned media or a set of angiogenic factors in the vessel network culture protocol. Our rationale stemmed from pilot experiments of fibroblasts and ECs in coculture, where we observed that over a 14-day period, fibroblasts consistently overgrew the vessel channel and migrated into the side channels, blocking perfusion of the vessel network and rendering the coculture unstable for long-term culture (Figure S2). Two FB:EC ratios were tested, with EC cell densities remaining constant while varying FB cell densities only such that effects could be attributed to FBs alone. To evaluate the importance of FB-EC crosstalk, we removed the FBs and tested ECs in monoculture with FB conditioned media, which was collected from FBs grown in 3D gel on-chip to better mimic our microfluidic coculture condition. For comparison, we also replaced the FBs with angiogenic factors and used EGM-2 as basal media with ECs in monoculture as a control. See Table 1 for a summary of these conditions.

To induce interstitial flow, we chose to use gravity-driven flow generated by differences in hydrostatic pressure in the media reservoirs (Figure 1f). Our pilot tests showed that flow could be maintained for 5 hours after establishing the initial hydrostatic pressure difference, after which the flow exponentially decayed and became static (Figure S3). Regarding the need for EC monolayers along the gel surface in the side channels, the monolayers were critical for our experiments as they (i) formed a physiological barrier that mitigated fibroblast outgrowth and undesired angiogenic sprouting; (ii) enabled robust anastomosis of the vessel network to the side channels; and (iii) ensured any perfusion induced by gravity-driven flow occurred through the vessel network. Without the monolayers, ECs and fibroblasts proliferated significantly outside the central gel compartment and gravity-driven flow caused leaked flow through the gel external to the vessels.

### Perfusability and leakage of MVNs

To assess perfusability as a primary function of our cultured MVNs, we performed flow visualization tests with fluorescent microbeads flowing through the network (Figure 2). Interestingly, not all vessel branches were perfused with microbeads even though they were luminal in structure (Figure 2c). These non-perfused branches were often long and tortuous, and thus high in flow resistance. Since microbeads travelled through the network along paths of least resistance, the high-resistance vessels were bypassed. We also observed that vessel walls appearing smooth and bright in fluorescence signal (Figure 2e) were better sealed and less prone to leakages, as opposed to vessel walls that appeared jagged and dim (Figure 2d and 2f), which often were more porous as evidenced by the leakage of microbeads into the interstitial space surrounding these types of vessels. Overall, MVN partial perfusion beads pattern can be summarized into 3 cases: 1) insufficient lumen formation, 2) non-perfusable region and 3) high resistance branches (Figure S4). To quantify perfusability and barrier integrity, we devised novel metrics where an MVN unit was considered perfusable if greater than 50% of the total vessel area was occupied by microbeads. Within each perfusable MVN unit, if beads were present in the interstitial space outside of the vessel area above a threshold area, the unit was considered “leaky”.

**Figure 2:**
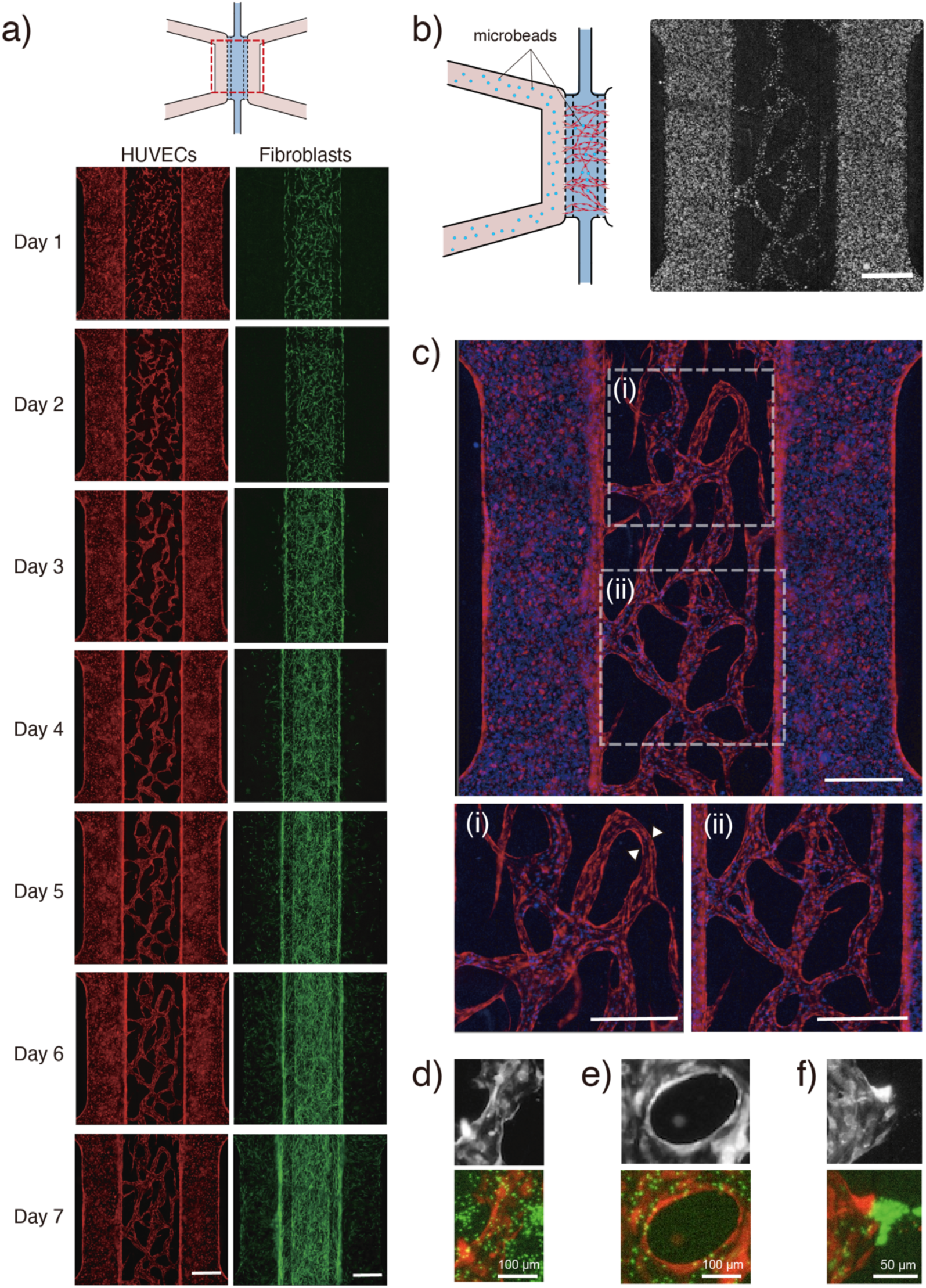
a) Fluorescence images showing vessel network development over 7 days (images shown are for FB1:EC5). Red: RFP-HUVECs; green: GFP-normal human lung fibroblasts. b) Illustration of fluorescent microbeads perfused through MVN and sample fluorescence image of microbeads in MVN. c) Overlay fluorescence image of microbeads (blue) in the vessel network (red), with two insets. (i) Magnification of inset showing a vessel branch with no beads, suggesting perfusion did not reach that branch (white arrows). (ii) Magnification of inset with all vessel branches perfused by beads. Scale bar = 500 mm for a) to c). d) Beads leaked out of vessels frequently near branches with jagged edges. e) Vessel branches with smooth edges contained the beads without leakage. f) Localized vessel leak near a vessel opening. (Top: raw image of vessels; Bottom: red = vessels, green = beads).

### Quantitative Morphological Metrics of MVNs

MVNs cultured with different experimental conditions (Table 1) showed qualitatively unique morphologies (Figure 4a). Both the FB1:EC5 and FB5:EC5 cocultures showed long and thin defined branches. Vessels in FB5:EC5 appeared denser than those in FB1:EC5 and showed significant vessel fusion over time. EC monocultures with FB conditioned media (FB-CM) did not appear to form any vessels and consisted of EC islands or patches separated by round void spaces. Vessel networks with angiogenic factors (Ang-V50) and with EGM-2 as control both exhibited a combination of defined vessels and EC patches, with vessel walls showing more jagged morphology than vessels in FB-EC coculture.

**Figure 3:**
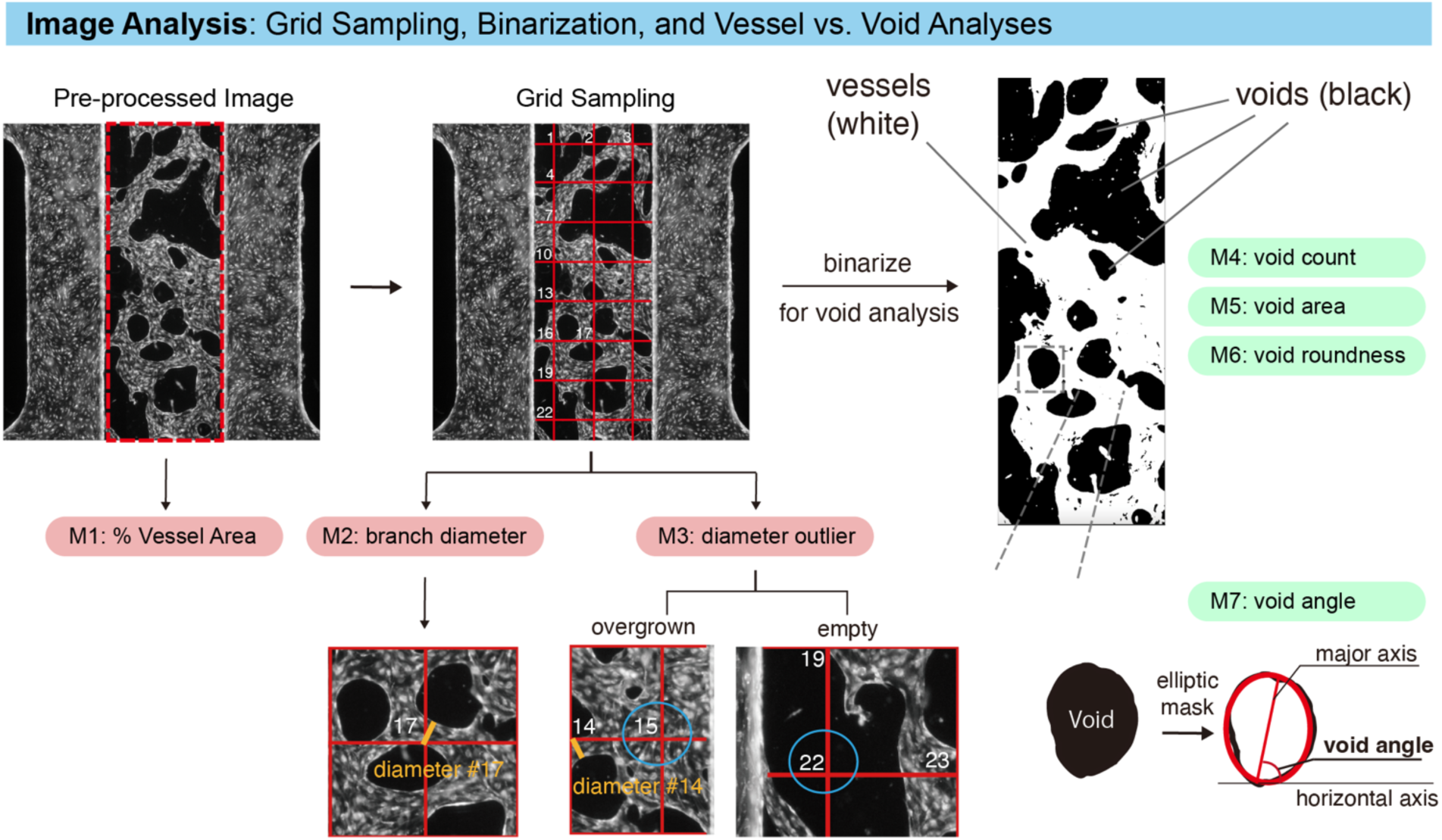
Image analysis: Grid sampling, binarization, and vessel vs. void analyses. Pre-processed fluorescence microscopy images were binarized for particle analysis to extract vessel-and void-based metrics (M1-M3 and M4-M7, respectively). Void angle was defined as the angle between the horizontal x-axis and the major axis of the elliptical fit of the void. Vessel diameters were measured at 24 points using systematic grid sampling. Outliers of diameter measurements were defined as grid points either with no vessels (grid point within a void) or within a large EC patch.

**Figure 4:**
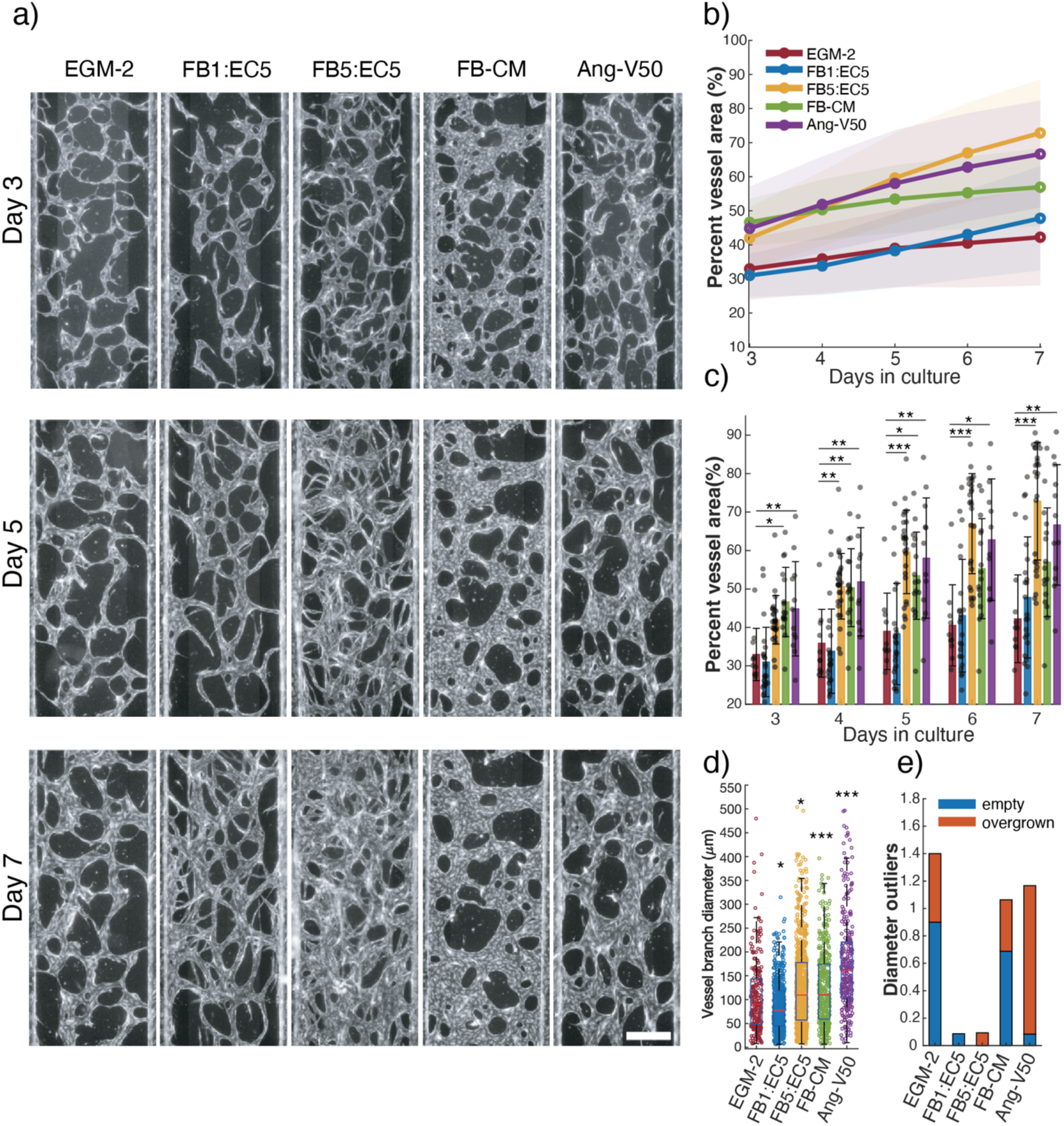
Morphometric analysis of fibroblast effects on MVN morphology: branched-based metrics. a) Fluorescence images of RFP-HUVECs forming MVNs in the middle channel. Scale bar = 500 mm. b) Line plots of average percent vessel area over time. Shaded regions represent standard deviations. c) Bar graph of average percent vessel area grouped by days in culture. Each data point represents percent vessel area of an individual device unit (n=10, 23, 26, 16, and 12 for EGM-2, FB1:EC5, FB5:EC5, FB-CM, and Ang-V50, respectively). d) Boxplot of individual vessel branch diameters (measurable, as defined in Figure 3) on Day 7. Red line is the median; box ranges from 25^th^ to 75^th^ percentiles. Each data point represents a single vessel diameter, collected from all the device units for each condition. e) Bar graph of number of diameter outliers (as defined in Figure 3) per device unit. For d) and e), n = 10, 23, 32, 16, and 12 for EGM-2, FB1:EC5, FB5:EC5, FB-CM, and Ang-V50, respectively. A few vessel units included in d) and e) were excluded from b) and c) because the prerequisite binary conversion using a global threshold failed to capture the true morphology of the vessel network in these units. For all graphs: *p<0.05, **p<0.01, ***p<0.001.

To further analyze vessel morphologies quantitatively, we created a list of seven morphology metrics (see Table 2). Because of the diverse array of complex morphologies observed, automated branch-focused image analysis programs (e.g., AngioTool, NIH) were unable to adequately describe the morphologies in our MVNs, prompting us to create our own list of metrics and custom image analysis scripts. For example, because we chose to include EC patches in our analysis to contrast island and vessel morphologies, it was necessary to create a custom method that could account for both islands and vessels in the same region of interest. We created a systematic grid-point sampling method (Figure 3), which had a sufficient sampling density to cover a significant portion of all vessel branches within a network while simultaneously providing a metric referred to as “diameter outliers”, which quantifies the frequency of EC islands (i.e., “overgrown”) or lack of vessels (i.e., “empty”) in the ROI.

**Table 2.** List of Seven Quantitative Morphological Metrics of Microvascular Networks.

| ***Metric*** | ***Description*** | ***Rationale*** |
| --- | --- | --- |
| Percent Vessel Area | Percent of ROI covered by vessels or EC patches | • Common metric in vessel network studies<br>• Indicator of overall vessel density without accounting for connectivity or alignment |
| Vessel Diameter | Diameter of individual vessels | • Common metric in vessel network studies<br>• Indicator of vessel size as assessment of the network's resemblance to vascular beds *in vivo* |
| Diameter Outliers | For each grid-point sampling that generates 24 points of interrogation in an ROI, this metric counts the number of instances where the grid point is within a void ("empty") or on an EC patch with no clear vessel width ("overgrown") | • A unique metric for this study<br>• An indicator of overall vessel network quality, as zero outliers suggests an optimal network morphology of evenly spaced defined vessel branches, no presence of EC patches, and minimal voids |
| Void Count | Number of void spaces enclosed by vessels; includes voids adjacent to ROI boundaries | • An alternative indicator for vessel density and vessel connectivity<br>• High void count means high vessel connectivity<br>• Low void count means either hyper-dense network with few voids, or lack of vessel connectivity leading to large void spaces |
| Average Void Roundness | Average (mean) of the roundness of individual voids; uses the same circularity metric as that used for particles | • An indicator of void shape and vessel branch pattern<br>• Circular voids (i.e., high roundness) tend to be surrounded by wide and short branches<br>• Elongated voids (i.e. low roundness) tend to be surrounded by long and thin branches |
| Average Void Area | Average (mean) of the area of individual voids | • In conjunction with void count, an indicator of vessel density |
| Void Angle | Angle of the major axis of a void from the horizontal *x*-axis; void is approximated as an ellipse using a particle analyzer | • An indicator of overall vessel alignment based on void angle distribution |

We further proposed novel void-based metrics (i.e., measurements describing the empty interstitial spaces between branches) that provided useful complementary information to traditional branch-based metrics, ultimately leading to new insights in MVN cultures. Table 2 provides a complete list of morphological metrics used in this study including their descriptions and rationales.

### Effects of Cocultured Fibroblasts on MVNs

In our first set of experiments, we compared the seven morphological metrics in Table 2 across five different culture conditions listed in Table 1 (i.e., FB1:EC5; FB5:EC5; FB-CM, Ang-V50, and EGM-2). Microscopy images were acquired daily between Days 3 to 7, allowing us to track temporal dynamics of MVN morphology by plotting each metric as a time course and examining both the metric and the rate of change of the metric based on the slope of the curve.

For Percent Vessel Area (PVA), FB5:EC5 had the highest average 5-day growth rate (14.7% per day), and the highest Day-3 PVA growth rate (20.7% from d3 to d4) (Figure 4b), suggesting that coculturing ECs with FBs at the higher cell density increased vessel area growth rate. FB-CM had high PVA on days 3 and 4 like FB5:EC5 (Figure 4c), but the growth rate drastically reduced on days 5 and 6, causing FB-CM to have the slowest average 5-day growth rate among all conditions (4.4% per day; statistically similar to the EGM-2 control at 5.6%) (Figure 4b). Ang-V50 had high PVA like FB5:EC5, and a medium average 5-day growth rate (9.7 % per day), suggesting that the angiogenic factor media composition can approach vessel area growth similar to cocultures with high FB cell density. Interestingly, FB1:EC5 produced a similar d3-d4 daily PVA as the EGM-2 control, but a higher average 5-day growth rate (FB1:EC5=10.8%, EGM-2=5.6%). Data appeared to show that low FB cell seeding density led to slow initial growth that eventually accelerated later in the time course.

Vessel diameters and diameter outliers together provide an indication of overall vessel network quality based on size uniformity of vessels and the frequency of non-vessel-like EC patches in the culture zone. We found that using vessel diameters alone was insufficient for assessing overall morphology, but that using vessel diameters in conjunction with diameter outliers uniquely distinguished vessel network types. Based on vessel diameter alone, Ang-V50 had the largest vessels, while FB5:EC5 and FB-CM had medium vessels that were smaller than Ang-V50 vessels but were larger than the vessels for FB1:EC5 and EGM-2 (Figure 4d). Analysis of diameter outliers, however, revealed that both the FB-EC cocultures have very few diameter outliers, exhibiting more defined measurable vessel branches that are evenly distributed across the culture zone as determined by our grid-point sampling method. Thus, although FB-CM and Ang-V50 produced networks with large branch diameters, they exhibited more EC patches and large void spaces, thus lowering the overall network quality. The EGM-2 control had small vessel diameters like FB1:EC5, but many diameter outliers (both “empty” and “overgrown”). These data are reflected in the fluorescence images (Figure 4a), where large void spaces and EC patches are more prevalent in EGM-2 than in FB1:EC5.

In addition to traditional branch-based metrics, we created novel void-based metrics that captured morphological behaviours not captured by branch-based metrics alone. Void count served as an alternative indicator of vessel density and connectivity, where a high void count means high vessel connectivity (i.e., a void surrounded by vessels) and a low void count means either a hyper-dense network with few voids (i.e., packed network), or lack of vessel connectivity leading to large void spaces (i.e., sparse network) (Figure 5a). For FB5:EC5, void count peaked on Day 5 and coincided with significant branch fusion observed for Day 5 and beyond, resulting in fewer voids (Figure 5a). For FB1:EC5, void count increased throughout the time course. This trend coincided with a gradual decrease in the average void area (Figure 5b). Taken together, these opposing trends for void count and void area matched our observations of consistent neovessel formation, where a lower number of initial large voids were split into a higher number of smaller voids over time due to a growing number of vessel branches. EGM-2, FB-CM and Ang-V50 all showed a similar trend of decreasing void counts and minimal change in void area over time, representing no significant neovessel formation after Day 3 (Figure 5a, 5b). Thus, while Ang-V50 and FB-CM supported initial vessel branch growth leading to high void counts on Day 3 (Figure 5d), these conditions failed to sustain vessel branch formation from Days 3 to 7 in a comparable manner to FB1:EC5.

**Figure 5:**
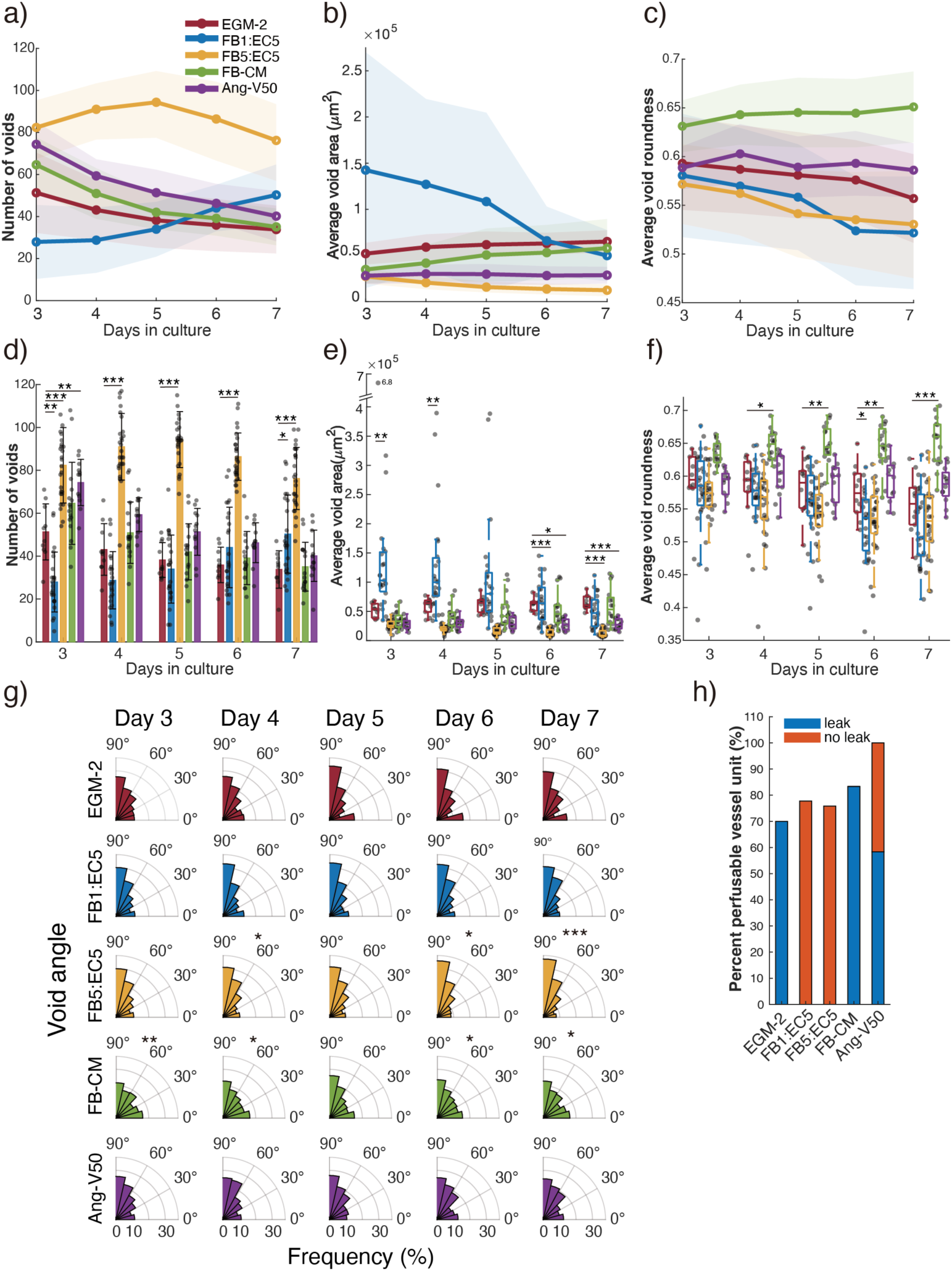
Morphometric analysis of fibroblast effects on MVN morphology: void-based metrics. a) Temporal line plots and d) bar graph (grouped by days in culture) of void count per device unit. b) Temporal line plots and e) box plot (grouped by days in culture) of average void area per device unit. c) Temporal line plots and f) box plot (grouped by days in culture) of average void roundness per device unit. Shaded regions represent standard deviations. Each data point represents either total void count in a device unit for d), or the average void area or void roundness across all voids in a device unit for e) and f). g) Polar histograms of void angles over time. Angles measured from the horizontal x-axis as defined in Figure 3. Angles between 90 to 180 degrees were converted to their supplementary angles. For a) to g): *n* = 10, 23, 26, 16, and 12 for EGM-2, FB1:EC5, FB5:EC5, FB-CM, and Ang-V50, respectively. h) Bar graph of percent perfusable vessel networks, calculated as number of perfusable MVN units divided by total tested MVN device units. Blue: perfusable units with leakage Red: perfusable units without leakage. For h): n = 10,18,29,12, and 12 for EGM-2, FB1:EC5, FB5:EC5, FB-CM, and Ang-V50, respectively. *p<0.05, **p<0.01, ***p<0.001.

Two other related void-based metrics were analyzed: (i) void roundness, representing the roundness? of individual voids (Figure 5c, 5f), and (ii) void angle, representing the angle between the major axis of the best-fit ellipse of the void and the horizontal *x*-axis (Figure 5g). Both metrics served as complementary measures of vessel alignment and stemmed from our observations that long defined vessels tended to extend in the longitudinal direction of the microchannels and were surrounded by elongated voids aligned in the same direction. We hypothesized that low void roundness values and void angles nearing 90 degrees correlated with MVNs displaying long defined vessels. Morphological analyses revealed that this was indeed the case. The two FB-EC cocultures had the smallest average void roundness measures (Figure 5d) due to the formation of long defined vessels interconnected by shorter vessel segments (Figure 4a). Furthermore, void roundness decreased over time in the FB-EC cocultures, with the most significant decreases coinciding with significant increases in void counts (Days 5-7 for FB1:EC5); Days 3-5 for FB5:EC5 (Figure 5a, 5c). The two FB-EC cocultures also exhibited the highest percentage of void angles between 75-90 degrees over the time course, as confirmed by observation of longitudinal vessel and void alignment (Figure S5). Taken together, these data suggest that the average void roundness of the MVN decreases with neovessel formation in concert with increased average void angle, increased void count and decreased void area. In contrast, FB-CM had the highest average void roundness, forming MVNs with distinct round void spaces that were not present in FB-EC cocultured networks. Although Ang-V50 and EGM-2 conditions appeared to have larger void roundness averages compared to cocultures, no significant differences were detected statistically. While FB-EC and Ang-V50 had relatively flatter distribution, Ang-V50 and FB-EC produced network with no directional alignment and co-culture networks demonstrated higher degree of alignment.

One interesting observation was that in MVNs with noticeable vessel alignment at the end on Day 7, the alignment was observed from the beginning on Day 1. However, not all MVNs with aligned vessels on Day 1 maintained the alignment through to Day 7. Thus, cultivating an initial environment that favours alignment is *necessary but not sufficient* to achieve successful alignment long-term. Based on this observation, and the fact that initial vessel alignment also occurs in non-cocultured conditions, we suspect that initial vessel alignment is driven more by the underlying fibrin gel structure rather than the presence of fibroblasts. Instead, fibroblasts helped maintain alignment later in the culture period, as cocultures tended to sustain the initial alignment while non-cocultures did not. Note that minor shifts in overall vessel alignment of the cultures may be due to the sprouting and growth of new transverse vessels that connect existing longitudinal vessels, which enhance network connectivity and perfusability, but may be manifested as decreases in void angle and increases in average void roundness.

Quantification of vessel leakage showed that MVN units in FB1:EC5 and FB5:EC5 had no leaks, while EC monoculture with EGM-2 media and with FB-CM exhibited leaks in all units (Figure 5h). EC monoculture with angiogenic factors (Ang-V50) had >50% of the units with leaks. Therefore, fibroblasts in MVN culture promoted higher EC wall integrity among the tested conditions. MVNs cultured with angiogenic factors had increased wall integrity compared to MVNs cultured in EGM-2 media or FB-CM, but not comparable to MVNs cultured with fibroblasts.

### Effects of VEGF concentration on MVNs

The moderate similarities we observed between Ang-V50 and FB5:EC5 prompted us to investigate whether the similarities could be further improved by simply modifying the composition or concentration of angiogenic factors, potentially allowing soluble factors to replace fibroblasts. Because VEGF is a critical factor in vessel development known to affect vessel diameter, fusion, and enlargement [57,58], we varied VEGF concentration while keeping the concentrations of FGF and S1P constant. To establish initial vessel development, we used the same amount of VEGF up to Day 3 and then switched to four different VEGF levels of 0, 10, 50 and 100 mg/mL from Day 3 onwards. Since complete EGM-2 media contains its own supplemented VEGF and FGF of unknown subtype, we removed the VEGF and FGF from the complete EGM-2 media to ensure the observed effects were only from the supplemented factors that we added with known subtype.

All VEGF concentrations tested exhibited EC island morphology (Figure 6a). VEGF100 had significantly smaller PVA and more void space compared to VEGF0 (Figure 6b, 6c), but no significant difference was observed between the VEGF concentrations for all other morphological metrics (Fig 6 and 7). Using microbeads to visualize MVN perfusability, we also found that all VEGF concentrations resulted in perfusable but leaky vessels (Figure 7h) The size of the leak, based on area coverage of microbeads outside the vessel, appeared similar among all VEGF concentrations, indicating no dependence of perfusability or endothelial barrier integrity on VEGF concentration.

**Figure 6:**
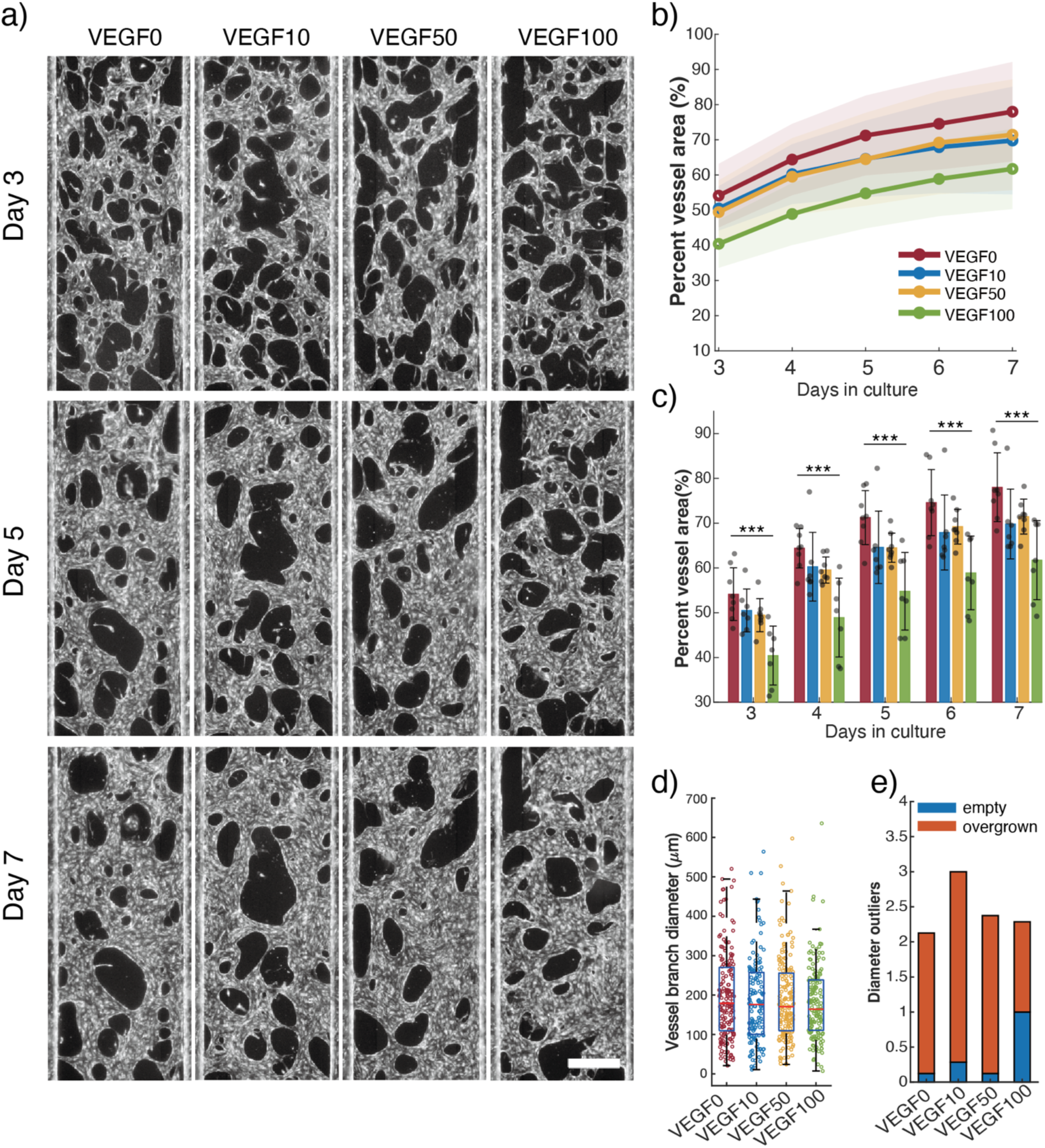
Morphometric analysis of VEGF concentration effects on MVN morphology: branched-based metrics. a) Fluorescence images of RFP-HUVECs forming MVNs in the middle channel. Scale bar = 500 mm. b) Line plots of average percent vessel area over time. Shaded regions represent standard deviations. c) Bar graph of average percent vessel area grouped by days in culture. Each data point represents percent vessel area of an individual device unit. d) Boxplot of individual vessel branch diameters (measurable, as defined in Figure 3) on Day 7. Red line is the median; box ranges from 25^th^ to 75^th^ percentiles. Each data point represents a single vessel diameter, collected from all the device units for each condition. e) Bar graph of number of diameter outliers (as defined in Figure 3) per device unit. *n* = 8, 7, 8, and 7 for VEGF0, VEGF10, VEGF50, and VEGF100, respectively. For all graphs: *p<0.05, **p<0.01, ***p<0.001.

**Figure 7:**
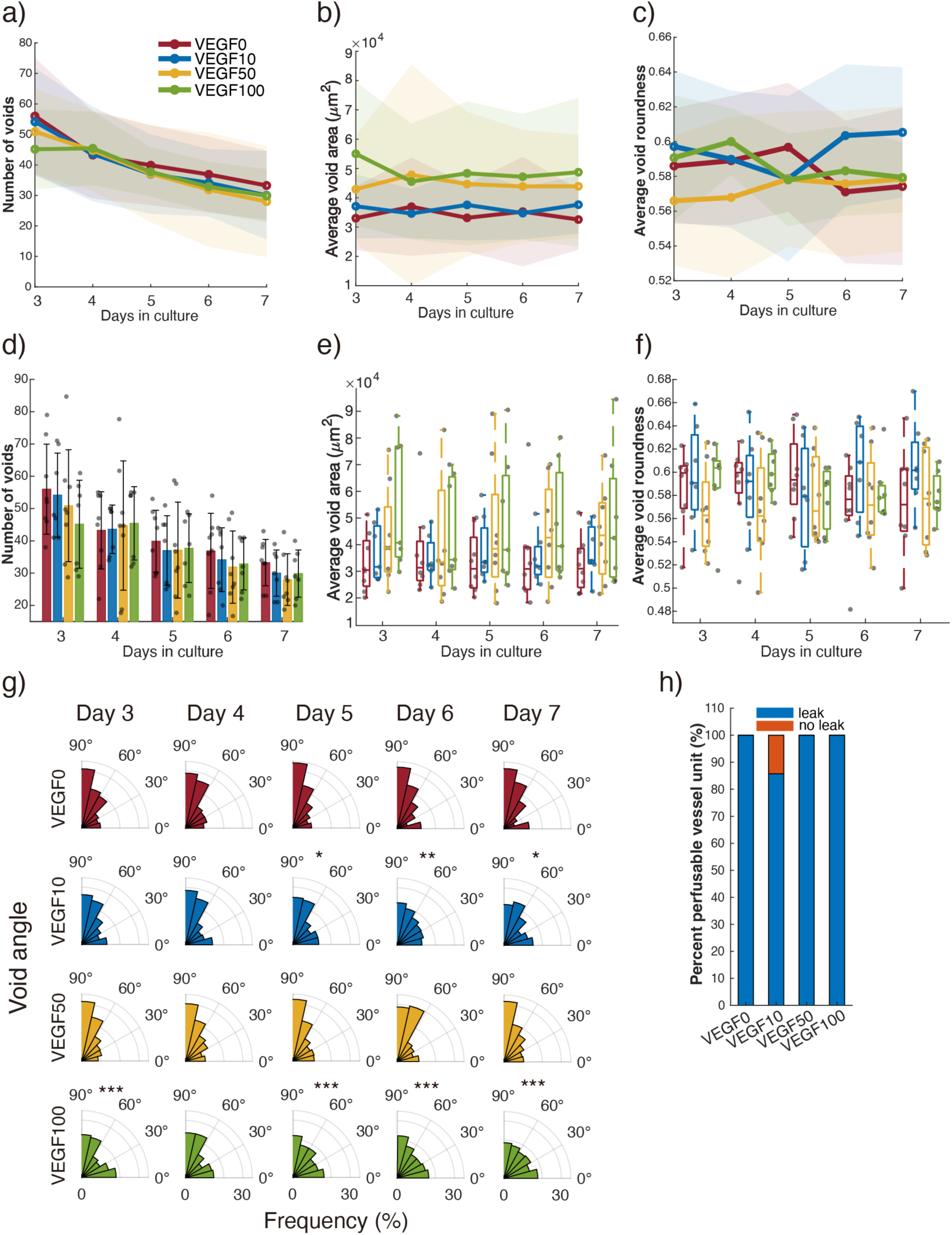
Morphometric analysis of VEGF concentration effects on MVN morphology: void-based metrics. a) Temporal line plots and d) bar graph (grouped by days in culture) of void count per device unit. b) Temporal line plots and e) box plot (grouped by days in culture) of average void area per device unit. c) Temporal line plots and f) box plot (grouped by days in culture) of average void roundness per device unit. Shaded regions represent standard deviations. Each data point represents either total void count in a device unit for d), or the average void area or void roundness across all voids in a device unit for e) and f). g) Polar histograms of void angles over time. Angles measured from the horizontal x-axis, as defined in Figure 3. Angles between 90 to 180 degrees were converted to their supplementary angles. h) Bar graph of percent perfusable vessel networks, calculated as number of perfusable MVN units divided by total tested MVN device units. Blue: perfusable units with leakage Red: perfusable units without leakage. *n* = 8, 7, 8, and 7 for VEGF0, VEGF10, VEGF50, and VEGF100, respectively. For all graphs: *p<0.05, **p<0.01, ***p<0.001.

Surprisingly, the morphology of VEGF50 MVNs (Figures 6 and 7) was qualitatively different than the morphology of Ang-V50 MVNs (Figures 4 and 5), even though they both contained the same concentration of supplemented VEGF (50 ng/ml). VEGF50 MVNs had a significantly lower void count and smaller average void area, but no other significant differences among the other metrics. We suspect that the 2 ng/ml VEGF and 2 ng/ml FGF that was removed from the EGM-2 media used in the VEGF50 condition led to these differences, as the withholding of those factors was the only difference between the two conditions.

### Effects of perfused flow on MVNs

Because vessel networks are known to remodel in the presence of flow [57], we tested the effect of flow on (i) FB-EC cocultured networks (FB1:EC5 with and without flow) and (ii) networks cultured with angiogenic factors (Ang-V20 with and without flow) (Figure 8). Images qualitatively showed that perfused flow reduced percent vessel area in FB1:EC5 but increased area coverage of EC islands in Ang-V20 (Figure 8a).

**Figure 8:**
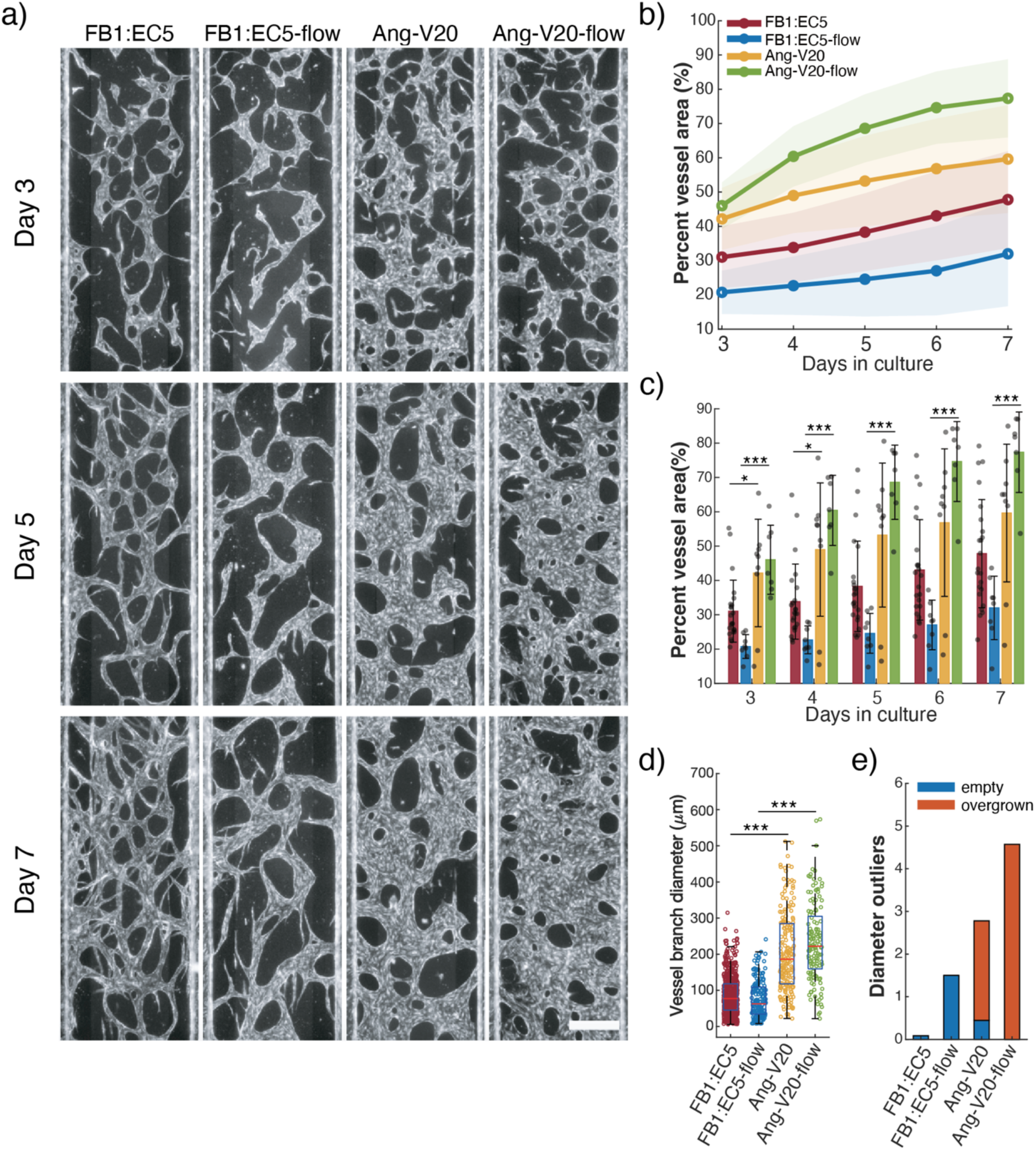
Morphometric analysis of perfusion flow effects on MVN morphology: branched-based metrics. a) Fluorescence images of RFP-HUVECs forming MVNs in the middle channel. Scale bar = 500 mm. b) Line plots of average percent vessel area over time. Shaded regions represent standard deviations. c) Bar graph of average percent vessel area grouped by days in culture. Each data point represents percent vessel area of an individual device unit. d) Boxplot of individual vessel branch diameters (measurable, as defined in Figure 3) on Day 7. Red line is the median; box ranges from 25^th^ to 75^th^ percentiles. Each data point represents a single vessel diameter, collected from all the device units for each condition. e) Bar graph of number of diameter outliers (as defined in Figure 3) per device unit. For all graphs: *n* = 23, 8, 9, and 7 for FB1:EC5, FB1:EC5-flow, Ang-V20 and Ang-V20-flow, respectively. *p<0.05, **p<0.01, ***p<0.001.

From quantitative morphometric analysis, we found that perfused flow led to: increased PVA growth rate for FB1:EC5 but decreased PVA growth rate for Ang-V20 (Figure 8b); reduced vessel diameter for FB1:EC5 but increased vessel diameter for Ang-V20 (Figure 8d, 8e); delayed branch formation for FB1:EC5 that began on Day 6 instead of Day 4 without flow (Figure 9a); and reduced void count but increased average void area for (Figure 9d and 9e). Altogether, these results suggest that FB1:EC5 with flow leads to reduced overall branch formation without effects on vessel diameters compared to the same coculture without flow, which is surprising given previous reports that flow increases vessel branch diameter[57]. The difference may be caused by the difference in device geometry and subsequent difference in mass transport pattern.

**Figure 9:**
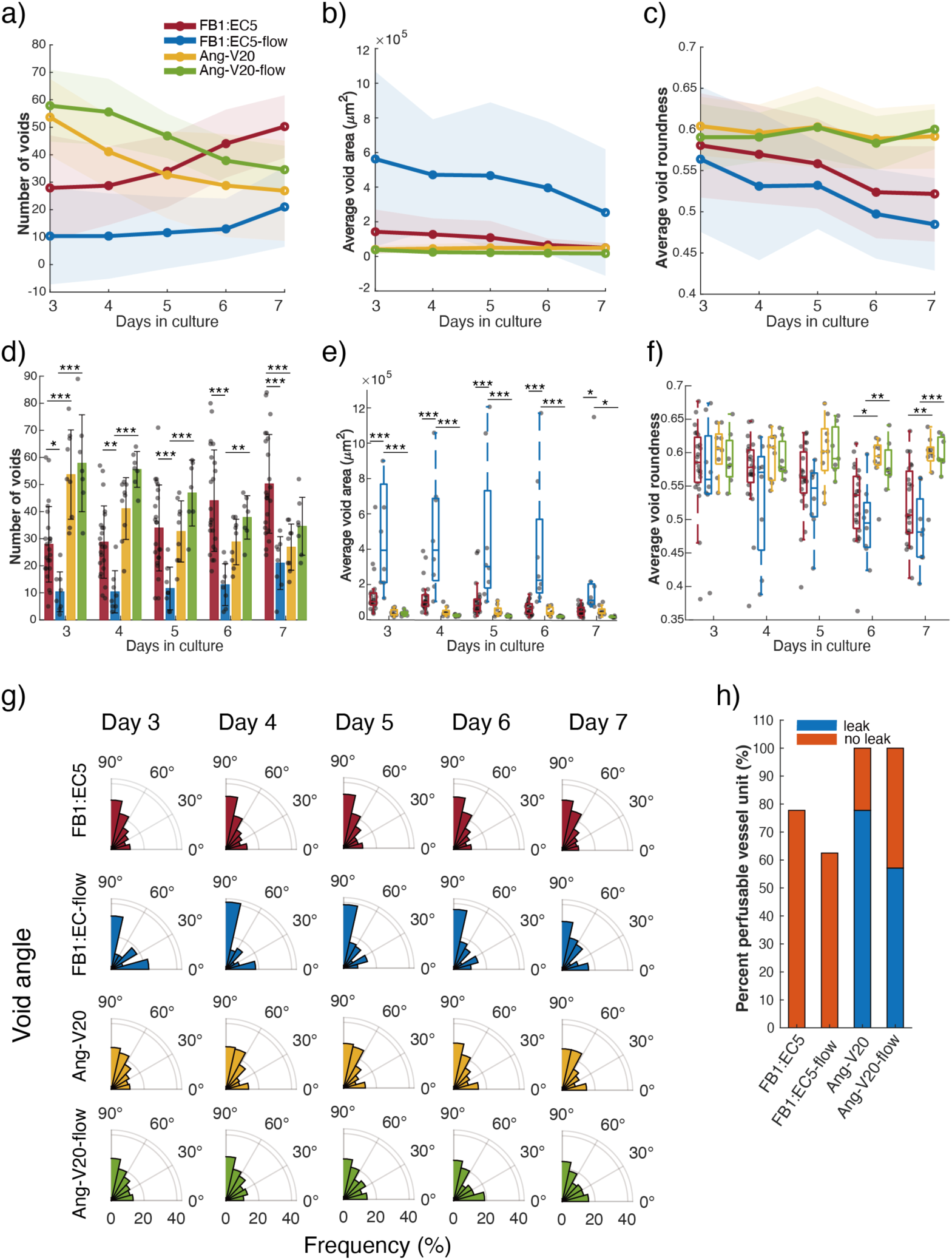
Morphometric analysis of perfusion flow effects on MVN morphology: void-based metrics. a) Temporal line plots and d) bar graph (grouped by days in culture) of void count per device unit. b) Temporal line plots and e) box plot (grouped by days in culture) of average void area per device unit. c) Temporal line plots and f) box plot (grouped by days in culture) of average void roundness per device unit. Shaded regions represent standard deviations. Each data point represents either total void count in a device unit for d), or the average void area or void roundness across all voids in a device unit for e) and f). g) Polar histograms of void angles over time. Angles measured from the horizontal x-axis, as defined in Figure 3. Angles between 90 to 180 degrees were converted to their supplementary angles. For a) to g): *n* = 23, 8, 9, and 7 for FB1:EC5, FB1:EC5-flow, Ang-V20, and Ang-V20-flow, respectively. h) Bar graph of percent perfusable vessel networks, calculated as number of perfusable MVN units divided by total tested MVN device units. Blue: perfusable units with leakage Red: perfusable units without leakage. For h): *n* = 18, 8, 9, and 7 for FB1:EC5, FB1:EC5-flow, Ang-V20, and Ang-V20-flow, respectively. *p<0.05, **p<0.01, ***p<0.001.

Using microbeads to visualize perfusability, vessel networks from both static and flow conditions were highly perfusable (Figure 9h), not including the cases with insufficient development of the vessel network that led to non-perfusable vessels. Like the other experiments in this study, leaky vessels were frequently observed in networks cultured without fibroblasts. Overall, flow did not have a major effect on overall perfusability in FB1:EC5 and Ang-V20 MVNs but may have increased endothelial barrier integrity in Ang-V20. Note that our morphometric analysis was applied to all cultured MVN units whether they were perfusable, and as such the analysis reflects the effect of our flow setup on vessel morphology rather than the effects of direct hemodynamic shear stress on the endothelium.

## Discussion

The goal of this study was to investigate how variations in culture conditions and experimental setups affect microvascular network (MVN) morphology in organ-on-a-chip (OOC) devices, with the aim of developing protocols that yield specific vascular features. We characterized MVN morphologies across a range of experimental conditions including with and without fibroblasts, conditioned media, angiogenic factors, and perfusion. In the process, we developed a comprehensive set of image-based quantitative metrics to describe MVN morphology, allowing us to distinguish MVNs into different morphology types.

In terms of selecting a device material for this study [59], we chose PDMS instead of thermoplastics, a decision that was informed by early experiments using acrylic-based microfluidic chips where we observed that vessel structures only formed near culture regions open to atmosphere (Figure S6). These early results suggested that sufficient gas diffusion was an essential factor in MVN formation. When we switched to gas-permeable PDMS, a significant difference was observed in MVN morphology and structure, leading us to choose PDMS for the rest of the study owing to its superior gas permeability. Note that plastic-based devices can still be used to create MVNs, as supported by our preliminary findings (Figure S6) and other published studies[60]. However, in our experience, plastic devices must employ either a more open-top design [61,62] (Figure S6) or rely on continuous media perfusion to deliver more oxygen to the cells.

In terms of microchannel compartmentalization, our MVN-on-a-chip used phaseguides[50,51], which allowed continuous, unobstructed connections and anastomoses between the vessel region and side channels. In contrast, micropost-based designs [63] require connections at predefined discrete locations, limiting adaptability but ensuring consistent geometry. A key advantage of the phaseguide approach is that perfusion can occur as long as a single connection to the endothelial lining in the side channels is established. This design also allows for the potential tuning of flow resistance, as the vessel network entrance is not constrained by the device geometry. Both approaches have trade-offs, and the choice should align with specific experimental goals.

To demonstrate network perfusability, we flowed fluorescent microbeads into one side channel and tracked the beads through the network. These bead perfusion tests showed that not all lumenized vessel branches were perfusable. Non-perfused branches found *in vivo* typically regress during vascular remodeling, resulting in optimized networks, but our *in vitro* self-assembled MVNs have not undergone this natural remodeling process. Even under flow conditions, we observed many non-perfused branches that persisted throughout the 7-day culture. Future work should explore extended culture or modified protocols to promote functional remodeling and increase the number of perfused branches.

We constructed temporal morphology curves based on daily vessel images to characterize dynamic changes in network morphology over time. This approach aims to emphasize the temporal dynamics of vessel development, as cultured vascular networks are living, evolving structures. A single static image is insufficient to determine whether a network will grow, regress, or undergo structural remodeling in subsequent days. By analyzing temporal profiles, researchers can select protocols tailored to specific rates of network development and accurately identify the current developmental stage on any given day.

To support our analysis, we introduced novel void-based metrics that address the wide variability in MVN morphologies that ranged from highly branched structures to patch-like formations. The void-based analysis offered a complementary perspective to traditional branch-based analysis, with strong potential for reconstructing global network architecture using simple features such as void area, roundness, orientation, and centroid position—parameters readily extracted using tools like the ImageJ Particle Analyzer. Given that the primary objective of this study was to evaluate protocol outcomes, we deliberately avoided imposing value judgments on network morphology or excluding samples with atypical structures. Instead, we presented a universally applicable analysis framework that combined branch-based and void-based metrics, enabling end users to select protocols best suited to their specific research goals and to define their own criteria for vascular quality.

Among the tested conditions, FB1:EC5 exhibited characteristics more closely resembling a physiologically healthy network in short-term culture, whereas FB5:EC5 demonstrated potential for rapid formation of dense networks and for investigating vessel fusion processes. Other conditions may be applicable for modeling pathological or leaky vasculature, or for culturing specific cell types while maintaining basic perfusion capacity. Future studies could benefit from extending the culture period of FB1:EC5, as the current 7-day timeline does not provide sufficient data to predict morphological trends beyond this point. It remains unknown whether FB1:EC5 exhibits a delayed fusion phase comparable to that observed in FB5:EC5. Expanding the range of profiled protocols may also facilitate the identification of a robust method for culturing quiescent, stable vascular networks that undergo minimal morphological changes upon reaching maturity.

Individually, the quantitative morphological metrics that were analyzed revealed various temporal patterns that appeared to correlate across metrics. These observed correlations prompted us to examine the temporal patterns in combination to find additional meta-patterns within the dataset that might not be immediately obvious. First, Table 3 summarizes the temporal trends for all experimental conditions across various morphological metrics, where an up-arrow indicates an upward trend, a down-arrow indicates a downward trend, and a dash indicates little or no change over time.

**Table 3:** Summary Table of the Trends in Void-based Metrics for All Conditions.

| Condition | Percent Vessel Area (PVA) | Void Count | Void Area |
| --- | --- | --- | --- |
| EGM-2 | ↑ | ↓ | ↑ |
| FB1:EC5 | ↑ | ↑ | ↓ |
| FB5:EC5 (Day 3-5) | ↑ | ↑ | ↓ |
| FB5:EC5 (Day 5-7) | ↑ | ↓ | ↓ |
| FB-CM | ↑ | ↓ | ↑ |
| Ang-V50 | ↑ | ↓ | — |
| FB1:EC5-flow | ↑ | ↑ | ↓ |
| Ang-V20 | ↑ | ↓ | — |
| Ang-V20- flow | ↑ | ↓ | ↓ |
| VEGF0 | ↑ | ↓ | — |
| VEGF10 | ↑ | ↓ | — |
| VEGF50 | ↑ | ↓ | — |
| VEGF100 | ↑ | ↓ | — |

↑ = metric increase with time, ↓ = decrease with time, — = slope of zero with small fluctuation

Examining Table 3 reveals three distinct groups of MVN morphological patterns: (i) UUD representing new branch formation; (ii) UDU representing vessel pruning; and (iii) UDD representing branch fusion (Figure 10). As an example, increasing percent vessel area (PVA) in combination with increasing void count and decreasing void area occurs when a new branch forms in the network, splitting one large void into multiple small voids. Conversely, decreasing void count in conjunction with increasing void area occurs when an existing branch is pruned, allowing multiple smaller voids to merge into a single larger void. When we compared our hypothetical “vessel events” illustrated in Figure 10 to the actual raw images acquired, we found remarkable correspondence between the two. Note that for several conditions related to VEGF, void area had neither an upward or downward trend over time. When matched with the raw images, we found that these cases corresponded with MVNs that exhibited comparable levels of vessel pruning and branch fusion that offset each other.

**Figure 10:**
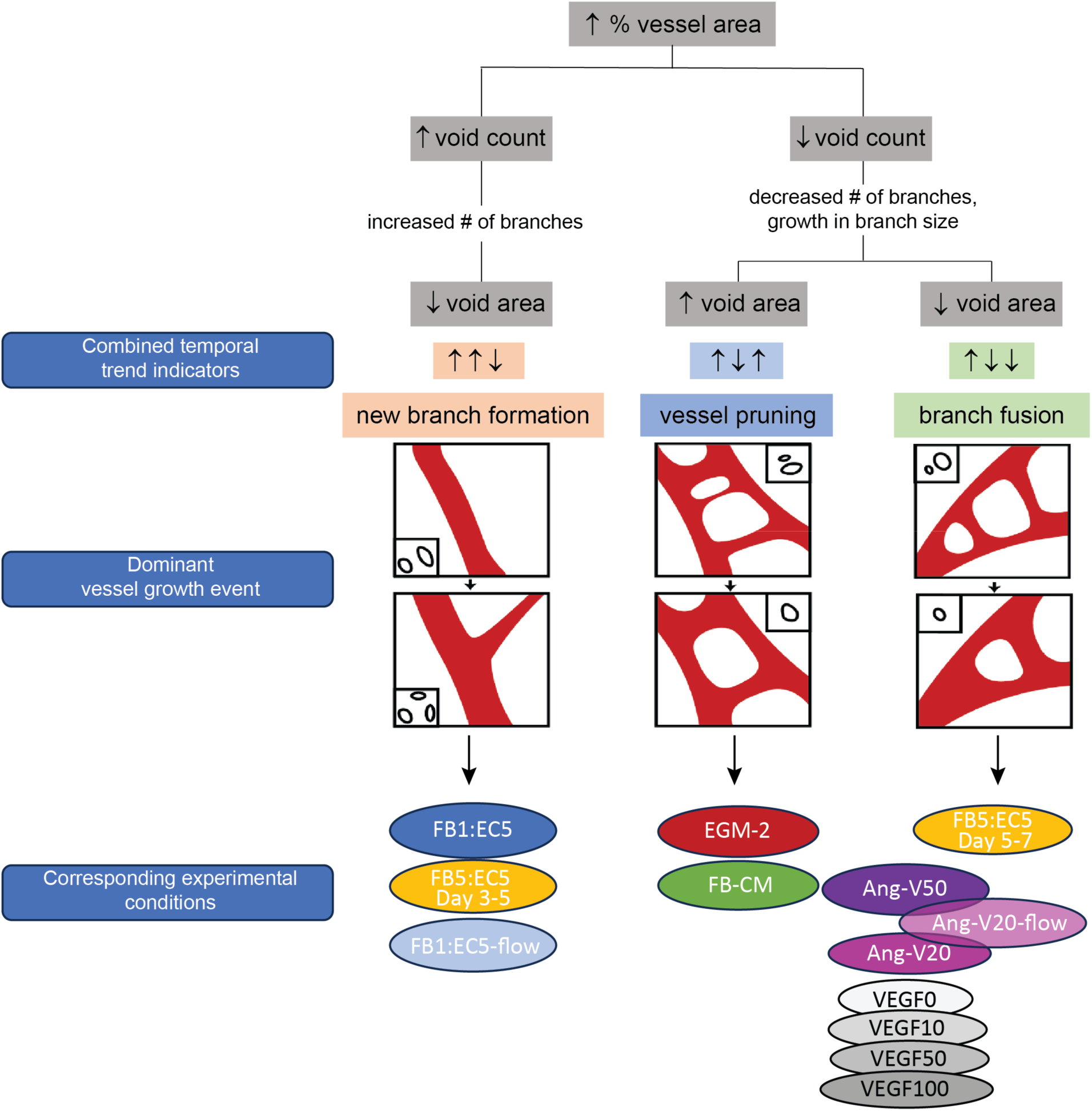
Tree diagram of various combined temporal trend indicators and their corresponding growth events and experimental conditions.

Another way to organize the data is to create 2D (or 3D) metric-based maps showing where the various experimental conditions are positioned in the 2D (or 3D) parameter space. As an example, we selected three relevant void-based metrics—void count, void area, and void shape— and plotted them in pairs (Figure 11). When characterizing a MVN by morphology, these metrics effectively capture the majority of the MVN’s most salient structural features. The plots highlight how similar various conditions are based on these metrics. Based on these plots, we observed that vessel morphology in FB1:EC5 and EGM-2 was similar across all three metrics, while Ang-V50 resembled FB5:EC5. FB-CM stood out due to its round void shape. The shaded boxes reflect temporal changes from Day 3 to Day 7, showing that the two FB-EC co-cultures exhibited the most significant morphological changes. In contrast, Ang-V50 and EGM-2 showed smaller changes relative to the others. In Figure 11, the four VEGF conditions clustered together near Ang-V50, suggesting that varying VEGF concentration does not significantly affect vessel morphology. We thus concluded that VEGF concentration was not a determining factor for vessel morphology in EC monoculture networks exposed to angiogenic factors. Overall, conditions with different VEGF levels did not produce morphology comparable to FB5:EC5.

**Figure 11:**
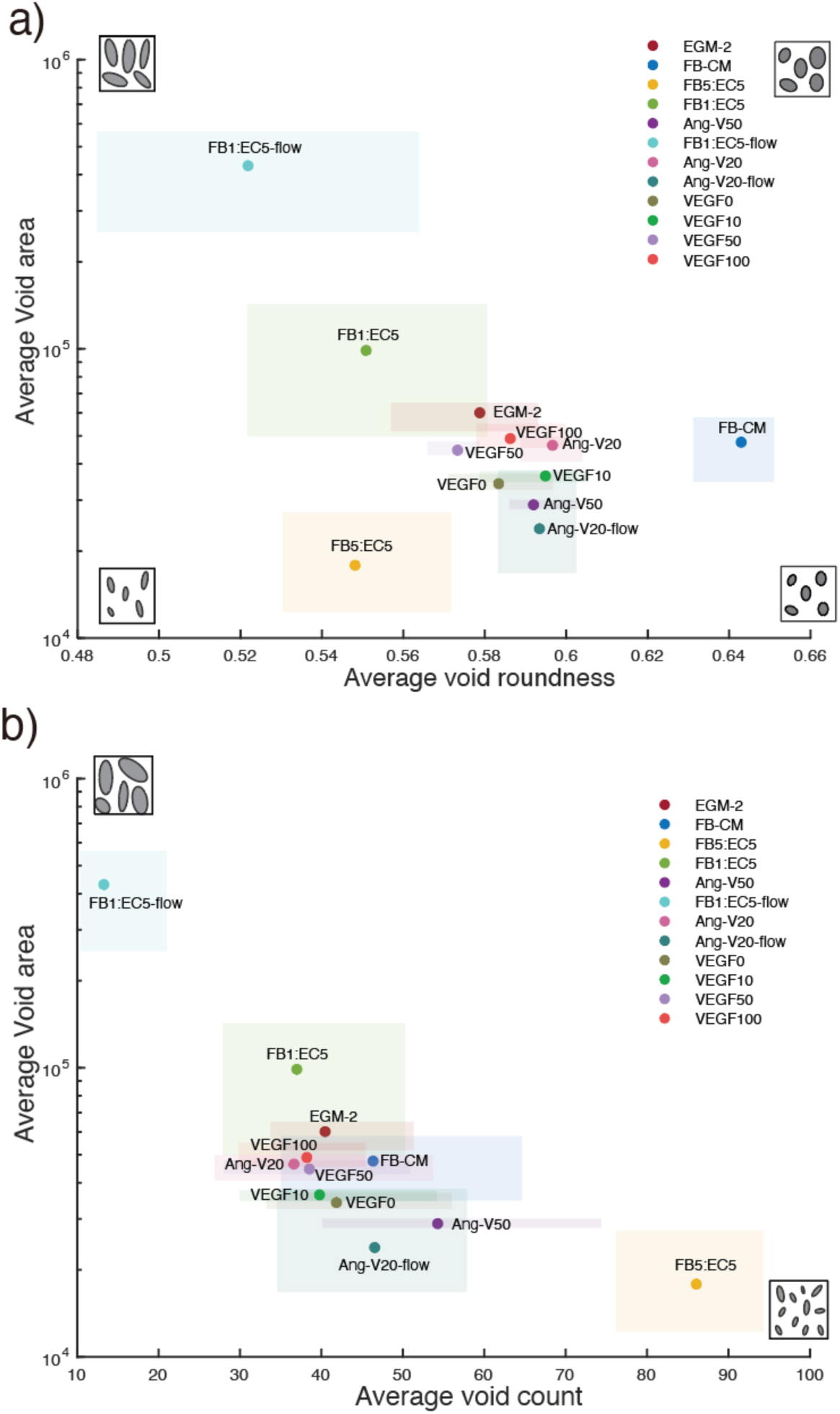
2D parameter mapping of experimental conditions. a) Average void roundness vs. average void area, and b) void count vs. average void area for all conditions. Y-axes are log scale. Each data point represents the average of the corresponding metric value between Days 3-7. The shaded box regions represent the maximum and minimum values between Day 3-7. Icons at the corners of each graph are illustrations of representative vessel morphology corresponding to that region of the map (grey areas represent voids and white spaces represent vessels in the icons).

Summarizing results from all metrics above, we found that: (a) the main morphology difference between the FB-EC co-cultures came from differences in branch diameter where high concentration of fibroblasts caused increase in growth rate, formation of new branches and branch outward growth; and (b) compared to FB-EC co-cultures, FB conditioned media or Ang-V50 could not replace cocultured fibroblasts and still result in the same morphology, though some similarities in the metrics were observed. Although EGM-2 had similar morphology to FB1:EC5 and reduced growth compared to FB5:EC5, EGM-2 MVNs also had major leakages and higher morphology variability among replicates indicating less consistency. Although the conditions without fibroblasts cannot be used to completely replace fibroblasts in vessel network culture, they presented unique morphology features that could make them useful for creating non-standard vessel networks for disease modelling and other applications.

We presented our efforts to quantify MVN morphology with a novel void-based approach. While our current analysis remains somewhat limited, additional features can potentially be extracted from the voids within the MVN. Future studies could explore more advanced techniques such as fractal analysis for deeper characterization, as well as lacunarity measures[64,65], both of which have been successfully applied in clinical vessel image quantification. A major advantage of the void-based method is its simplicity -vessel morphology can be described with just four metrics: void location, area, roundness, and void angle. In contrast, branch-based methods require more complex data and are less effective for irregular MVNs. This compact set of features is also well-suited for machine learning applications, enabling dimensionality reduction for tasks such as image reconstruction and automated image analysis.

## Conclusion

In this study, we systematically evaluated the impact of fibroblast presence, soluble factors, and luminal flow on MVN morphology, perfusability, and vessel wall integrity using a microfluidic vasculature-on-a-chip platform. Our results highlight the essential role of direct fibroblast incorporation in promoting physiologically relevant, thin, and interconnected vascular networks, an effect that cannot be replicated by soluble cues alone. We also introduced a novel void-based morphological analysis that sensitively captures differences in vessel-free space characteristics with fewer parameters than traditional branch-based methods. This simplified, high-content feature set is well-suited for integration with machine learning and automated image analysis, offering new opportunities for dimensionality reduction and predictive modeling. Future work will expand this void-based approach to include advanced metrics such as fractal dimension and lacunarity, enabling deeper characterization of MVN architecture and further enhancing its utility in vascular tissue engineering and organ-on-a-chip applications.

## Acknowledgements

The authors have no conflict of interest to disclose. We acknowledge the financial support from Canada Foundation for Innovation (CFI) John R Evans Leadership Fund (#32171), Natural Sciences and Engineering Research Council of Canada (NSERC) Discovery Grant (RGPIN-2019-5885), and the Early Researcher Award from Ontario Ministry of Research and Innovation (ER15-11-058). The authors gratefully acknowledge technical support from the Centre for Research and Applications in Fluidic Technologies (CRAFT). Some figures were created with BioRender.

## References

1. Novosel EC, Kleinhans C, Kluger PJ. Vascularization is the key challenge in tissue engineering. Adv Drug Deliv Rev. 2011 Apr 30;63(4):300–11.

2. Leung CM, de Haan P, Ronaldson-Bouchard K, Kim GA, Ko J, Rho HS, et al. A guide to the organ-on-a-chip. Nat Rev Methods Primer. 2022 May 12;2(1):1–29.

3. Ingber DE. Human organs-on-chips for disease modelling, drug development and personalized medicine. Nat Rev Genet. 2022 Aug;23(8):467–91.

4. Vunjak-Novakovic G, Ronaldson-Bouchard K, Radisic M. Organs-on-a-chip models for biological research. Cell. 2021 Sep 2;184(18):4597–611.

5. Jain A, Barrile R, van der Meer AD, Mammoto A, Mammoto T, De Ceunynck K, et al. Primary Human Lung Alveolus-on-a-chip Model of Intravascular Thrombosis for Assessment of Therapeutics. Clin Pharmacol Ther. 2018 Feb;103(2):332–40.

6. Jeon JS, Bersini S, Whisler JA, Chen MB, Dubini G, Charest JL, et al. Generation of 3D functional microvascular networks with human mesenchymal stem cells in microfluidic systems. Integr Biol Quant Biosci Nano Macro. 2014 May;6(5):555–63.

7. Soon K, Li M, Wu R, Zhou A, Khosraviani N, Turner WD, et al. A human model of arteriovenous malformation (AVM)-on-a-chip reproduces key disease hallmarks and enables drug testing in perfused human vessel networks. Biomaterials. 2022 Sep;288:121729.

8. Campisi M, Shin Y, Osaki T, Hajal C, Chiono V, Kamm RD. 3D self-organized microvascular model of the human blood-brain barrier with endothelial cells, pericytes and astrocytes. Biomaterials. 2018 Oct;180:117–29.

9. Huh D, Leslie DC, Matthews BD, Fraser JP, Jurek S, Hamilton GA, et al. A human disease model of drug toxicity-induced pulmonary edema in a lung-on-a-chip microdevice. Sci Transl Med. 2012 Nov 7;4(159):159ra147.

10. Wang P, Jin L, Zhang M, Wu Y, Duan Z, Guo Y, et al. Blood–brain barrier injury and neuroinflammation induced by SARS-CoV-2 in a lung–brain microphysiological system. Nat Biomed Eng. 2024 Aug;8(8):1053–68.

11. Choi JH, Choi HK, Lee KB. In Situ Detection of Neuroinflammation using Multi-cellular 3D Neurovascular Unit-on-a-Chip. Adv Funct Mater. 2023 Nov 9;33(46):2304382.

12. Walji N, Kheiri S, Young EWK. Angiogenic Sprouting Dynamics Mediated by Endothelial-Fibroblast Interactions in Microfluidic Systems. Adv Biol. 2021;5(11):2101080.

13. Song JW, Bazou D, Munn LL. Anastomosis of endothelial sprouts forms new vessels in a tissue analogue of angiogenesis. Integr Biol Quant Biosci Nano Macro. 2012 Aug;4(8):857–62.

14. Nagaraju S, Truong D, Mouneimne G, Nikkhah M. Microfluidic Tumor-Vascular Model to Study Breast Cancer Cell Invasion and Intravasation. Adv Healthc Mater. 2018 May;7(9):e1701257.

15. Chen MB, Whisler JA, Jeon JS, Kamm RD. Mechanisms of tumor cell extravasation in an in vitro microvascular network platform. Integr Biol Quant Biosci Nano Macro. 2013 Oct;5(10):1262–71.

16. Zhao Y, Wu Y, Islam K, Paul R, Zhou Y, Qin X, et al. Microphysiologically Engineered Vessel-Tumor Model to Investigate Vascular Transport Dynamics of Immune Cells. ACS Appl Mater Interfaces. 2024 May 8;16(18):22839–49.

17. Staples SCR, Yin H, Sutherland FSK, Prescott EK, Tinney D, Hamilton DW, et al. Intussusceptive angiogenesis-on-a-chip: Evidence for transluminal vascular bridging by endothelial delamination. Proc Natl Acad Sci U S A. 2025 Apr 22;122(16):e2423700122.

18. Lee S, Ko J, Park D, Lee SR, Chung M, Lee Y, et al. Microfluidic-based vascularized microphysiological systems. Lab Chip. 2018 Sep 11;18(18):2686–709.

19. Mandrycky CJ, Howard CC, Rayner SG, Shin YJ, Zheng Y. Organ-on-a-chip systems for vascular biology. J Mol Cell Cardiol. 2021 Oct;159:1–13.

20. Zhang S, Wan Z, Kamm RD. Vascularized organoids on a chip: strategies for engineering organoids with functional vasculature. Lab Chip. 2021 Feb 9;21(3):473–88.

21. Huang CBX, Tu TY. Recent advances in vascularized tumor-on-a-chip. Front Oncol. 2023;13:1150332.

22. Wang YI, Abaci HE, Shuler ML. Microfluidic blood–brain barrier model provides in vivo-like barrier properties for drug permeability screening. Biotechnol Bioeng. 2017;114(1):184–94.

23. Barrile R, van der Meer AD, Park H, Fraser JP, Simic D, Teng F, et al. Organ-on-Chip Recapitulates Thrombosis Induced by an anti-CD154 Monoclonal Antibody: Translational Potential of Advanced Microengineered Systems. Clin Pharmacol Ther. 2018;104(6):1240– 8.

24. van Dijk CGM, Brandt MM, Poulis N, Anten J, van der Moolen M, Kramer L, et al. A new microfluidic model that allows monitoring of complex vascular structures and cell interactions in a 3D biological matrix. Lab Chip. 2020 May 19;20(10):1827–44.

25. Offeddu GS, Possenti L, Loessberg-Zahl JT, Zunino P, Roberts J, Han X, et al. Application of Transmural Flow Across In Vitro Microvasculature Enables Direct Sampling of Interstitial Therapeutic Molecule Distribution. Small Weinh Bergstr Ger. 2019 Nov;15(46):e1902393.

26. Guarino V, Perrone E, De Luca E, Rainer A, Cesaria M, Zizzari A, et al. Pericyte-Assisted Vascular Lumen Organization in a Novel Dynamic Human Blood-Brain Barrier-on-Chip Model. Adv Healthc Mater. 2025 May 6;e2401804.

27. Bischel LL, Young EWK, Mader BR, Beebe DJ. Tubeless microfluidic angiogenesis assay with three-dimensional endothelial-lined microvessels. Biomaterials. 2013 Feb;34(5):1471– 7.

28. Liu Y, Sakolish C, Chen Z, Phan DTT, Bender RHF, Hughes CCW, et al. Human in vitro vascularized micro-organ and micro-tumor models are reproducible organ-on-a-chip platforms for studies of anticancer drugs. Toxicology. 2020 Dec 1;445:152601.

29. Tsvirkun D, Grichine A, Duperray A, Misbah C, Bureau L. Microvasculature on a chip: study of the Endothelial Surface Layer and the flow structure of Red Blood Cells. Sci Rep. 2017 Mar 24;7:45036.

30. Watanabe M, Salvadori A, Markovic M, Sudo R, Ovsianikov A. Advanced liver-on-chip model mimicking hepatic lobule with continuous microvascular network via high-definition laser patterning. Mater Today Bio. 2025 Jun;32:101643.

31. Zhang B, Montgomery M, Chamberlain MD, Ogawa S, Korolj A, Pahnke A, et al. Biodegradable scaffold with built-in vasculature for organ-on-a-chip engineering and direct surgical anastomosis. Nat Mater. 2016 Jun;15(6):669–78.

32. Russell T, Dirar Q, Li Y, Chiang C, Laskowitz DT, Yun Y. Cortical spheroid on perfusable microvascular network in a microfluidic device. PloS One. 2023;18(10):e0288025.

33. Quintard C, Tubbs E, Jonsson G, Jiao J, Wang J, Werschler N, et al. A microfluidic platform integrating functional vascularized organoids-on-chip. Nat Commun. 2024 Feb 16;15(1):1452.

34. Wan Z, Zhong AX, Zhang S, Pavlou G, Coughlin MF, Shelton SE, et al. A Robust Method for Perfusable Microvascular Network Formation In Vitro. Small Methods. 2022;6(6):2200143.

35. Jeon JS, Bersini S, Gilardi M, Dubini G, Charest JL, Moretti M, et al. Human 3D vascularized organotypic microfluidic assays to study breast cancer cell extravasation. Proc Natl Acad Sci U S A. 2015 Jan 6;112(1):214–9.

36. Vila Cuenca M, Cochrane A, van den Hil FE, de Vries AAF, Lesnik Oberstein SAJ, Mummery CL, et al. Engineered 3D vessel-on-chip using hiPSC-derived endothelial- and vascular smooth muscle cells. Stem Cell Rep. 2021 Sep 14;16(9):2159–68.

37. Nahon DM, Vila Cuenca M, van den Hil FE, Hu M, de Korte T, Frimat JP, et al. Self-assembling 3D vessel-on-chip model with hiPSC-derived astrocytes. Stem Cell Rep. 2024 Jul 9;19(7):946–56.

38. Arslan U, van den Hil FE, Mummery CL, Orlova V. Generation and Characterization of hiPSC-Derived Vascularized-, Perfusable Cardiac Microtissues-on-Chip. Curr Protoc. 2024;4(7):e1097.

39. Maurissen TL, Spielmann AJ, Schellenberg G, Bickle M, Vieira JR, Lai SY, et al. Modeling early pathophysiological phenotypes of diabetic retinopathy in a human inner blood-retinal barrier-on-a-chip. Nat Commun. 2024 Feb 14;15(1):1372.

40. Hsu YH, Yang WC, Chen YT, Lin CY, Yang CF, Liu WW, et al. Spatially controlled diffusion range of tumor-associated angiogenic factors to develop a tumor model using a microfluidic resistive circuit. Lab Chip. 2024 May 14;24(10):2644–57.

41. Bi Y, Shirure VS, Liu R, Cunningham C, Ding L, Meacham JM, et al. Tumor-on-a-chip platform to interrogate the role of macrophages in tumor progression. Integr Biol Quant Biosci Nano Macro. 2020 Sep 30;12(9):221–32.

42. Zeinali S, Bichsel CA, Hobi N, Funke M, Marti TM, Schmid RA, et al. Human microvasculature-on-a chip: anti-neovasculogenic effect of nintedanib in vitro. Angiogenesis. 2018 Nov;21(4):861–71.

43. Moon BU, Li K, Malic L, Morton K, Shao H, Banh L, et al. Reversible bonding in thermoplastic elastomer microfluidic platforms for harvestable 3D microvessel networks. Lab Chip. 2024 Oct 22;24(21):4948–61.

44. Vempati P, Popel AS, Mac Gabhann F. Extracellular regulation of VEGF: isoforms, proteolysis, and vascular patterning. Cytokine Growth Factor Rev. 2014 Feb;25(1):1–19.

45. Phng LK, Belting HG. Endothelial cell mechanics and blood flow forces in vascular morphogenesis. Semin Cell Dev Biol. 2021 Dec;120:32–43.

46. Winkelman MA, Kim DY, Kakarla S, Grath A, Silvia N, Dai G. Interstitial flow enhances the formation, connectivity, and function of 3D brain microvascular networks generated within a microfluidic device. Lab Chip. 2021 Dec 21;22(1):170–92.

47. Tronolone JJ, Jain A. Engineering New Microvascular Networks On-Chip: Ingredients, Assembly, and Best Practices. Adv Funct Mater. 2021 Apr;31(14):2007199.

48. Nakamura M, Ninomiya Y, Nishikata K, Futai N. On-chip long-term perfusable microvascular network culture. Jpn J Appl Phys. 2022 May;61(SD):SD1040.

49. Zudaire E, Gambardella L, Kurcz C, Vermeren S. A Computational Tool for Quantitative Analysis of Vascular Networks. PLOS ONE. 2011 Nov 16;6(11):e27385.

50. Vulto P, Podszun S, Meyer P, Hermann C, Manz A, Urban GA. Phaseguides: a paradigm shift in microfluidic priming and emptying. Lab Chip. 2011 May 7;11(9):1596–602.

51. Mohan MD, Young EWK. TANDEM: biomicrofluidic systems with transverse and normal diffusional environments for multidirectional signaling. Lab Chip. 2021 Oct 26;21(21):4081–94.

52. Wong JF, Mohan MD, Young EWK, Simmons CA. Integrated electrochemical measurement of endothelial permeability in a 3D hydrogel-based microfluidic vascular model. Biosens Bioelectron. 2020 Jan 1;147:111757.

53. Wang X, Phan DTT, George SC, Hughes CCW, Lee AP. 3D Anastomosed Microvascular Network Model with Living Capillary Networks and Endothelial Cell-Lined Microfluidic Channels. Methods Mol Biol Clifton NJ. 2017;1612:325–44.

54. Devadas D, Moore TA, Walji N, Young EWK. A microfluidic mammary gland coculture model using parallel 3D lumens for studying epithelial-endothelial migration in breast cancer. Biomicrofluidics. 2019 Nov;13(6):064122.

55. Ogilvie IRG, Sieben VJ, Floquet CFA, Zmijan R, Mowlem MC, Morgan H. Reduction of surface roughness for optical quality microfluidic devices in PMMA and COC. J Micromechanics Microengineering. 2010 May;20(6):065016.

56. Guckenberger DJ, Groot TE de, Wan AMD, Beebe DJ, Young EWK. Micromilling: a method for ultra-rapid prototyping of plastic microfluidic devices. Lab Chip. 2015 May 20;15(11):2364–78.

57. Sano H, Watanabe M, Yamashita T, Tanishita K, Sudo R. Control of vessel diameters mediated by flow-induced outward vascular remodeling in vitro. Biofabrication. 2020 Jul;12(4):045008.

58. Drake CJ, Little CD. VEGF and vascular fusion: implications for normal and pathological vessels. J Histochem Cytochem Off J Histochem Soc. 1999 Nov;47(11):1351–6.

59. Berthier E, Young EWK, Beebe D. Engineers are from PDMS-land, Biologists are from Polystyrenia. Lab Chip. 2012 Mar 7;12(7):1224–37.

60. Song J, Choi H, Koh SK, Park D, Yu J, Kang H, et al. High-Throughput 3D In Vitro Tumor Vasculature Model for Real-Time Monitoring of Immune Cell Infiltration and Cytotoxicity. Front Immunol. 2021;12:733317.

61. Park S, Young EWK. E-FLOAT: Extractable Floating Liquid Gel-Based Organ-on-a-Chip for Airway Tissue Modeling under Airflow. Adv Mater Technol. 2021;6(12):2100828.

62. Newton JD, Song Y, Park S, Kanagarajah KR, Wong AP, Young EWK. Tunable In Situ Synthesis of Ultrathin Extracellular Matrix-Derived Membranes in Organ-on-a-Chip Devices. Adv Healthc Mater. 2024 Aug;13(20):e2401158.

63. Kim S, Lee H, Chung M, Jeon NL. Engineering of functional, perfusable 3D microvascular networks on a chip. Lab Chip. 2013 Mar 19;13(8):1489–500.

64. Di Ieva A, Grizzi F, Gaetani P, Goglia U, Tschabitscher M, Mortini P, et al. Euclidean and fractal geometry of microvascular networks in normal and neoplastic pituitary tissue. Neurosurg Rev. 2008 Jul;31(3):271–81.

65. Gould DJ, Vadakkan TJ, Poché RA, Dickinson ME. Multifractal and Lacunarity Analysis of Microvascular Morphology and Remodeling. Microcirc N Y N 1994. 2011 Feb;18(2):136–51.

