## Supplemental Materials for "Mechanical, Biochemical, and Multicellular Effects on Vessel Network Morphometrics in a Microfluidic Vasculature-on-a-Chip"

#### Supplementary Methods

*Cell Culture Experiment Timelines*

Figure S1 illustrates the experimental timelines for cell seeding, media changes, sample collection, readouts, and other interventions for all MVN culture experiments conducted in this study. Detailed procedures are in the main text. These schematics are included for additional clarity.

*Image Analyses and Definitions of Metrics*

<u>*Pre-processing*</u>: All raw images went through contrast enhancement using histogram equalization. Day 3 to Day 7 images of the device were converted to a stack, aligned in reference to Day 7 image using the "*template matching"* plugin. The common vessel area in the middle channel across all 5 images in the stack was manually "*cropped*" and used for all analysis. The grey scale image stacks were converted to binary using the "*triangle*" thresholding method. "*Remove outlier*" was performed for both black and white pixels with radius=2 and threshold = 50.

<u>*Percent vessel coverage area*</u>: represents the percentage of vessel pixels (white) over the entire region of interest (ROI), measured with "*area fraction"* in the "*measurement"* function.

<u>***Vessel diameter:***</u> A grid with 24 points was laid over the pre-processed image stack. Vessel diameters were manually measured at these 24 grid points, with certain exceptions described below. Diameters were defined as the length of a line drawn perpendicular to the vessel branch passing

through each grid point. If no vessel branch passed directly through a grid point, the closest branch intersecting any of the four grid lines surrounding that point was measured instead. In these cases, the diameter was measured as the length of a line drawn perpendicular to the vessel branch at the intersection between the vessel and the grid line.

Occasionally, a "diameter" measured according to this method could exceed the length of the vessel branch itself. This suggests that the measurement location was too close to a node, making the diameter questionable as a true cross-sectional measurement. Therefore, vessel diameters were only recorded when the measured value was shorter than the length of the corresponding vessel branch. In cases where the diameter exceeded the branch length, the measurement line was repositioned to the nearest appropriate branch around the grid point, ensuring that the new diameter was shorter than the branch length.

***Measurement Outliers:*** A grid point was marked as an "N/A empty outlier" if no vessel branches passed through either the grid point or its adjacent grid lines, or if the closest vessel branch had already been measured at a previous grid point. This designation indicated the absence of a measurable vessel branch at that location. For typical vessel branches, the diameter measurement line was drawn from one vessel wall to the opposite wall. However, in some instances, the vessel structure was so large that the opposite wall extended into the next grid square or beyond. These measurements were labeled as "N/A overgrown outliers," indicating that the vessel sheet was too large to allow for a proper diameter measurement.

***Void Count, Void Area, Void Roundness, and Angle:*** The "*Analyze Particles*" function was used to highlight empty void spaces between vessel branches in the binary vessel images, with a *threshold* range set to 500–infinity. Void count was recorded as the number of highlighted regions. Void roundness represented the geometric roundness of these empty spaces. Void angle measured the orientation of these spaces, providing insight into the alignment of surrounding vessel branches. Void area represented the actual size of the empty regions.

For angle measurements, all voids were approximated as “*Ellipses*” using the “Analyze Particles” function. The void angle was defined as the angle between the major axis of each ellipse and the horizontal x-axis, calculated automatically using the "*Analyze Particles*" function. Angles between 90° and 180° were converted to their supplementary angles to maintain consistent orientation representation.

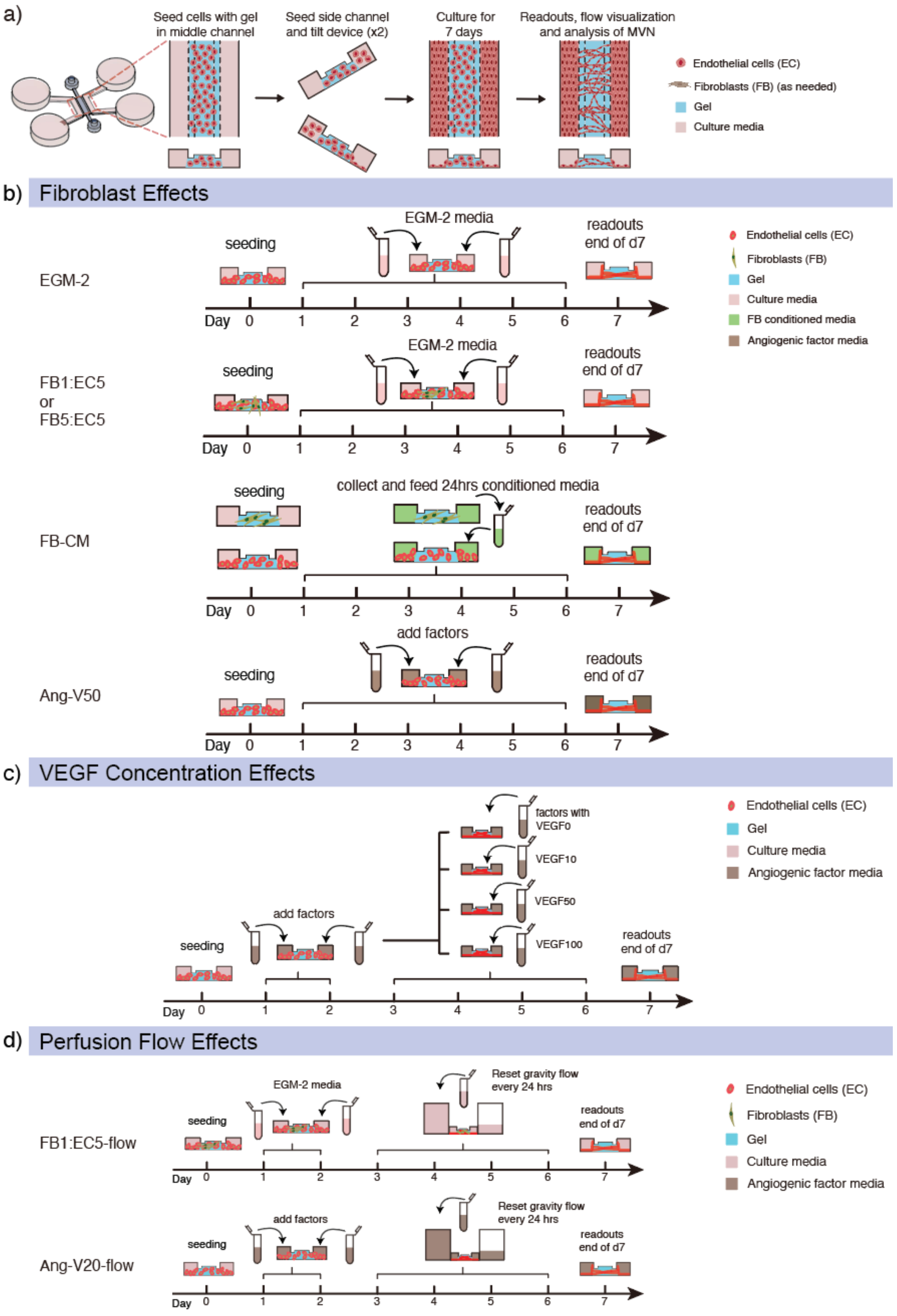
a)
Seed cells with gel in middle channel
Seed side channel and tilt device (x2)
Culture for 7 days
Readouts, flow visualization and analysis of MVN
Endothelial cells (EC)
Fibroblasts (FB) (as needed)
Gel
Culture media
b) Fibroblast Effects
EGM-2
seeding
EGM-2 media
readouts end of d7
Day 0 1 2 3 4 5 6 7
Endothelial cells (EC)
Fibroblasts (FB)
Gel
Culture media
FB conditioned media
Angiogenic factor media
FB1:EC5 or FB5:EC5
FB-CM
collect and feed 24hrs conditioned media
Ang-V50
add factors
c) VEGF Concentration Effects
factors with VEGF0
VEGF10
VEGF50
VEGF100
d) Perfusion Flow Effects
FB1:EC5-flow
Reset gravity flow every 24 hrs
Ang-V20-flow

**Figure S1**: MVN-on-a-chip cell culture procedure and schematics of the experimental timelines for cell seeding, media changes, sample collection, and readouts for all MVN culture experiments conducted in this study: a) Cells (ECs only or ECs and FBs) suspended in hydrogel were first seeded into the vessel channel. After polymerization of the gel, ECs were seeded into one side channel and then tilted to allow the ECS to attach to the wall of the gel. The other side channel was seeded and tilted in a similar fashion. The culture was maintained for 7 days based on the desired experimental conditions. After 7 days, the experiment was stopped, and readouts, flow visualization, and imaging were performed. (b) comparing effects of fibroblasts to fibroblast conditioned media (FB-CM) and angiogenic factors; (c) comparing different concentrations of VEGF; and (d) comparing effects of perfusion using gravity-driven flow.

### Other Observations

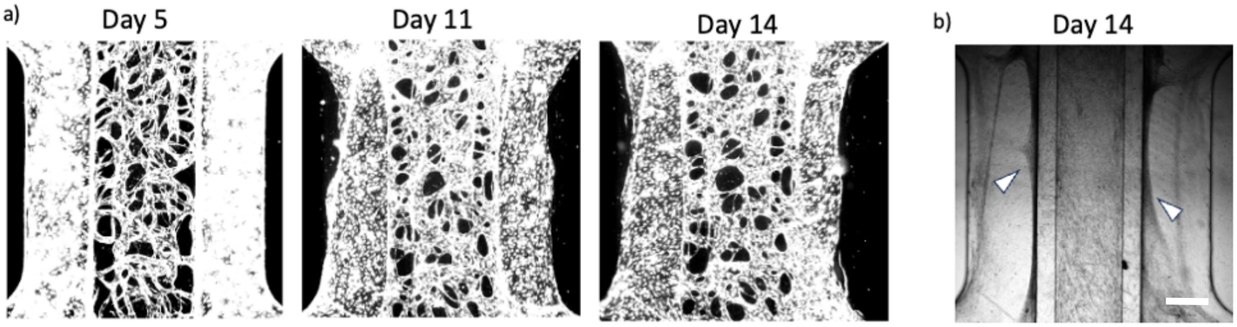


**Figure S2**: Vessel network fusion and fibroblast overgrowth in 14-day co-culture of ECs and FBs. a) Fluorescence images of vessel network (EC5:FB5) showing excessive branch fusion after Days 5, 11, and 14. b) Brightfield image of the vessel network co-culture. Dark shaded regions in the two side channels (white arrows) are overgrown fibroblasts, which block vessel network perfusion.

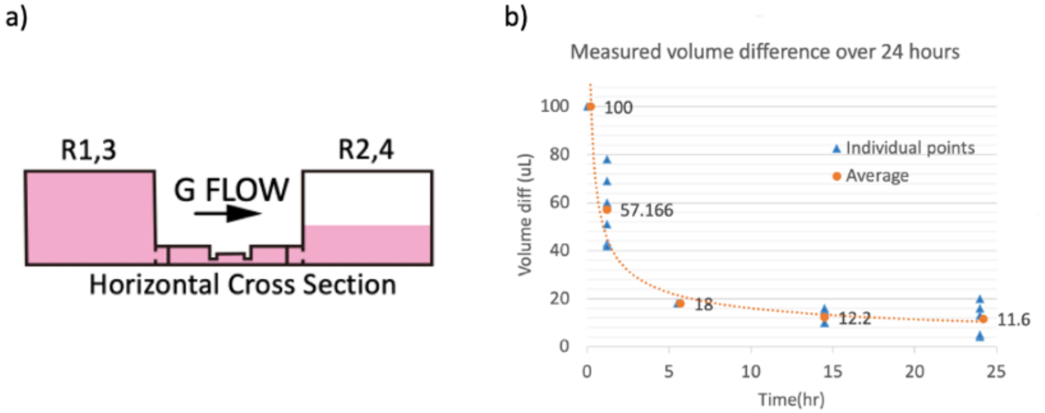


**Figure S3**: 24-hour gravity-driven flow profile. a) Illustration of gravity flow setup and measurement. 100 µL media was added to the two reservoirs on the left side (R1, R3) to allow perfusion to the right. b) Measured volume difference between total media volume of the left two reservoirs and right two reservoirs over 24-hour flow period. 100 µL media was added to the left two reservoirs at $t$ = 0 hr. Media volume in the left two reservoirs and right two reservoirs was measured with pipettes and the calculated difference was plotted to indicate presence of significant hydrostatic pressure difference and gravity-driven flow. After 5 hours of flow, the volume difference reached a plateau.

***Cases of Partially Perfusable MVNs***

Partial perfusion patterns of a microvascular network (MVN) fell into three distinct cases. In the first case, non-perfused vessels were narrow and likely did not form lumens (Figure S4a,d). When such narrow vessels were present near the junction between the vessel network and the side channels, all downstream branches (i.e., a large portion of the network) remained unperfused. In the second case, the perfusion pattern showed evidence of path selection, where beads preferentially traveled between nodes through branches that were shorter or straighter (Figure S4b,e). The longer or more tortuous branches, although appearing wide enough to have developed lumens and to allow perfusion, remained unperfused, likely due to their geometry causing higher fluidic resistance. This type of path-selective partial perfusion occurred more frequently in networks with high branch density or with a limited number of connections between the network and the side channels, both of which increase the number of possible paths between nodes. The third pattern showed a localized non-perfused region (Figure S4c,f), with a clear boundary between perfused and non-perfused areas. The non-perfused branches appeared wide enough but may not have been sufficiently tall in the z-axis to permit perfusion.

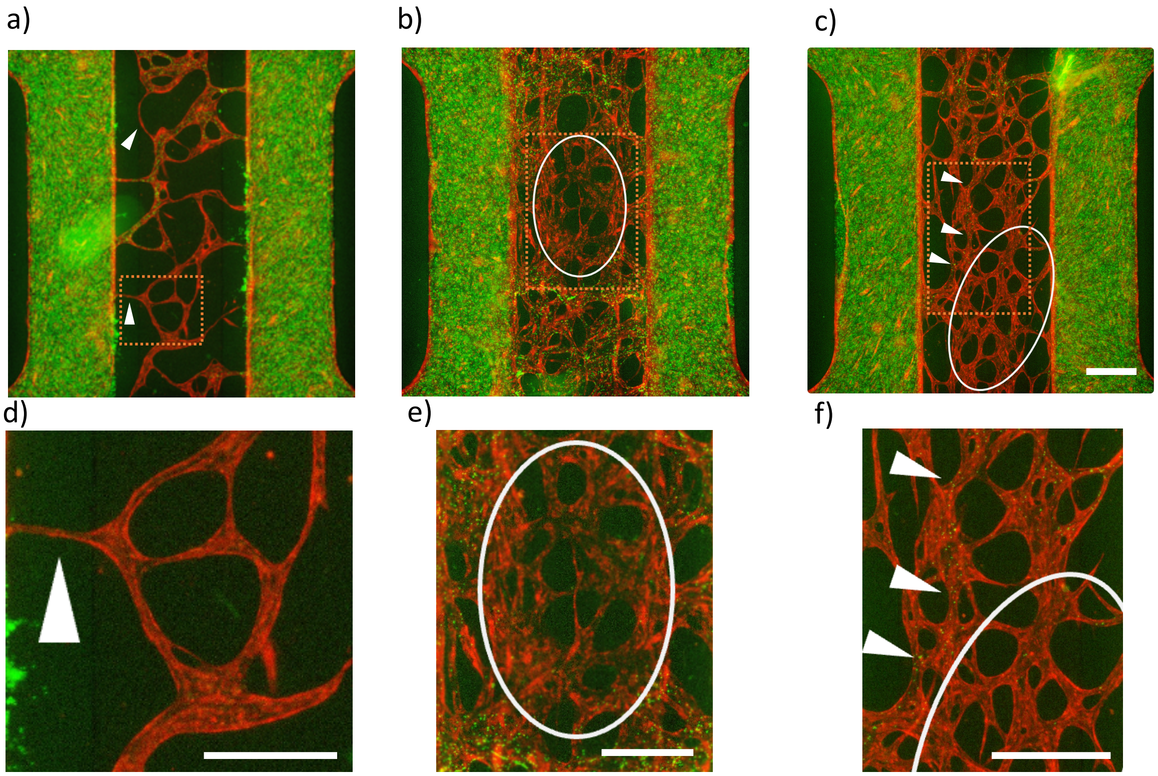


**Figure S4:** Microbead vessel perfusion pattern for partially perfused MVN. Red=vessel network. Green=microbeads. a) Image of partially perfused vessel network with insufficient lumen formation, and d) magnified region (inset in (a)) with arrow points to a branch with likely no lumen formation. b) Image of partially perfused vessel network with non-perfusable region (white oval) and e) magnified region (inset in (b)) of the non-perfusable region. c) Image of partially perfused vessel network with non-perfused branches of high resistance and f) magnified region (inset in (c)). Arrows point to low resistance perfused branch. White oval shows high resistance non-perfused branches.

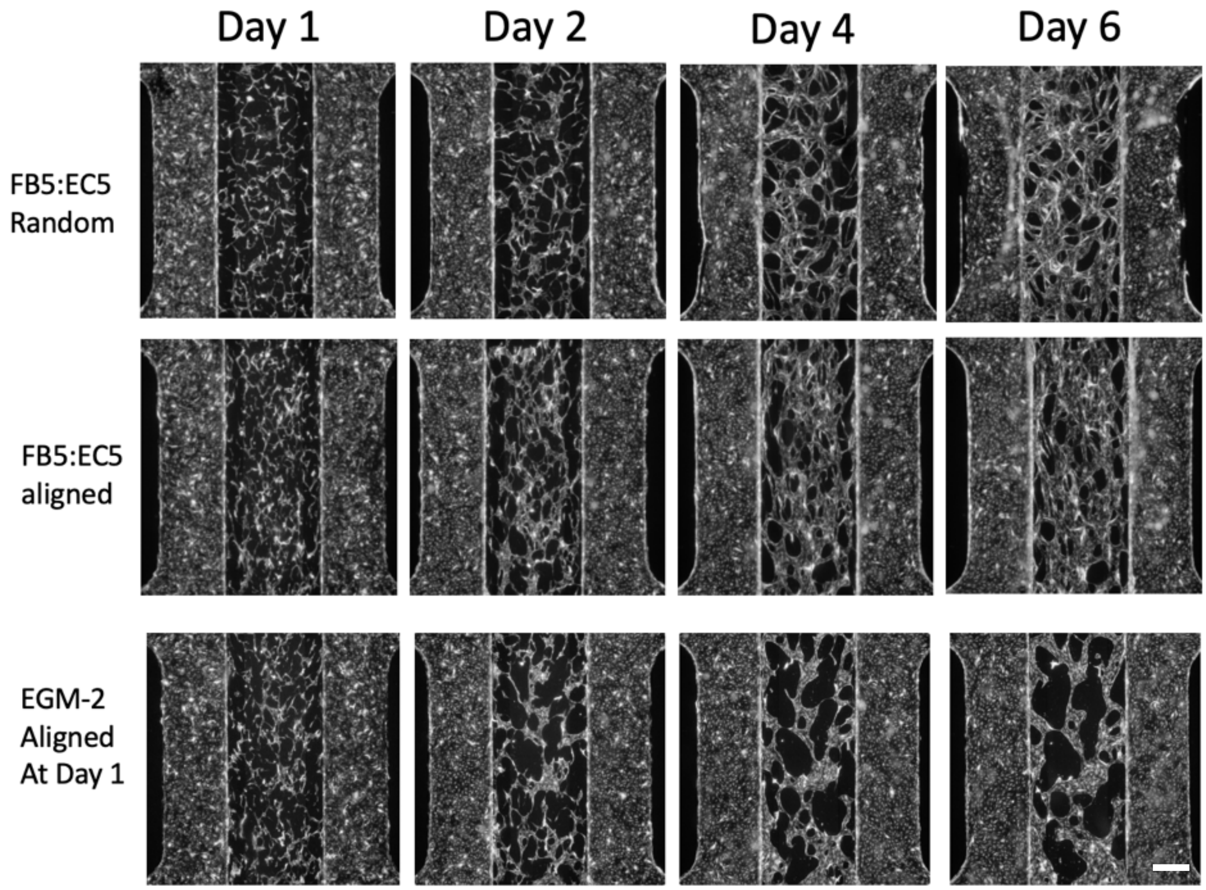


**Figure S5:** MVN with vessel alignment. Top row: fluorescence images of FB-EC coculture vessel network with no visible branch alignment. Middle row: fluorescent images of FB-EC coculture vessel network with visible branch alignment. Bottom row: fluorescent image of EC monoculture vessel network with visible branch alignment on Day 1 but loss of vessel alignment later in the culture.

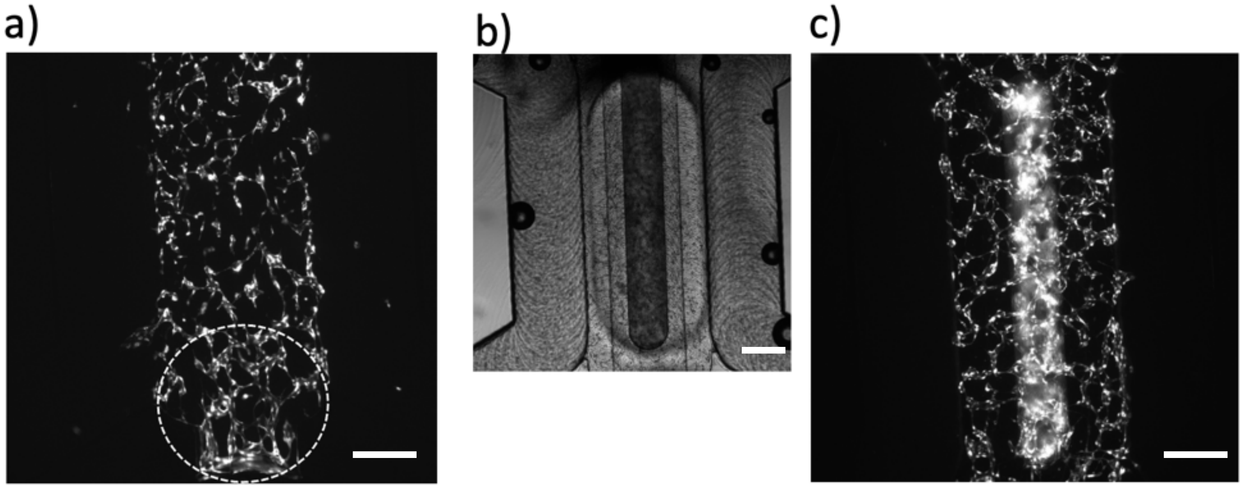


**Figure S6:** MVN culture in plastic devices. a) Fluorescence images of MVN cultured in a PMMA device that had the same geometry as the PDMS devices illustrated in Figure 1 of the main text. Connected vessel branches were only formed near channel reservoirs, which are exposed to air and allow for greater oxygen availability. b) Brightfield image of a PMMA device with open top for oxygen access of the MVN gel and c) the cultured MVN. Connected branches were successfully formed along the channel in the open top PMMA device, providing evidence of the need for high oxygen access for MVN culture.
